# Membrane-active peptides escape drug-resistance in cancer

**DOI:** 10.1101/2022.10.27.513961

**Authors:** Aurélie H. Benfield, Felicitas Vernen, Reuben S.E. Young, Ferran Nadal-Bufí, Heinz Hammerlindl, David J. Craik, Helmut Schaider, Nicole Lawrence, Stephen J. Blanksby, Sónia Troeira Henriques

## Abstract

Acquired drug-resistance is a recurring problem in cancer treatment, and this is particularly true for patients with metastatic melanoma that carry a BRAF V600E mutation. In the current study, we explored the use of membrane-active peptides as an alternative therapeutic modality to target drug-resistant melanoma cells. We produced slow-cycling and drug-resistant melanoma cells using dabrafenib, a small molecule drug that targets tumor cells with BRAF V600E mutation, and characterised their lipidome and proteome to investigate the role of membrane lipids in acquired drug-resistance. Despite some changes in the lipid composition, tested anti-melanoma membrane-active cyclic peptides (cTI and cGm) killed melanoma cells that are sensitive, tolerant, or resistant to dabrafenib. Importantly, melanoma cells did not develop resistance to cTI or cGm, nor changed their lipid composition with long-term peptide treatment. Therefore, these peptides are well suited as templates to design therapeutic leads to target drug-resistant metastatic melanoma cells and/or as co-treatment with small molecule drugs.

## Introduction

Melanoma is the deadliest and most aggressive form of skin cancer due to its metastatic and invasive potential. Current standards of care to treat metastatic melanoma patients include immunotherapy with antibodies to inhibit immune check-points (e.g., pembrolizumab and nivolumab) [1], and targeted therapy with small molecule drug kinase inhibitors to inhibit MAPK/ERK signalling (e.g., dabrafenib [2], which inhibits the mutant BRAF V600E occurring in 40-45% of metastatic melanoma). Melanoma patients that respond to immunotherapy have an increased survival rate; however, half of the patients are intrinsically resistant to immunotherapy. Patients that carry MAPK/ERK mutations respond well to targeted therapy with small molecule BRAF/MEK inhibitors, but continued therapy with these kinase inhibitors converts drug-sensitive into non-responsive tumors [3-5], which rapidly acquire drug resistance, enabling the disease to progress within six to ten months of treatment [6, 7]. Patients with metastatic melanoma that are resistant to current immunotherapy and targeted therapy require novel therapies with alternative mechanisms of actions.

Tumors that become non-responsive to BRAF/MEK inhibitors normally emerge from populations of slow-cycling cells that are tolerant to treatment with kinase inhibitors. The slow-cycling cells that survive drug treatment are referred to as drug-tolerant persisters (DTPs) [8, 3, 9], and reflect a non-mutational response to drug exposure. However, continued drug treatment of DTPs leads to a state of phenotypic diversity and further non-reversible permanent resistance [3, 10]. Therefore, new therapeutic strategies that prevent tumour heterogeneity, and eradicate populations of DTPs, are crucial to prevent acquired drug resistance [10, 9, 7].

It has been hypothesised that host-defense peptides (HDPs) can bypass established mechanisms of drug resistance and kill drug-tolerant and drug-resistant cancer cells [11-14]. These peptides selectively target the negatively charged membrane surface of cancerous cells, over neutral cell surfaces of healthy cells, and induce cell death by disruption of cell membranes, or by crossing cell membranes and acting on intracellular targets [15, 16, 14, 17]. Owing to their membrane-dependent mechanism of action, it has been hypothesised that HDPs can kill cancer cells that are slow-growing, cancer cells that are refractory to treatment with conventional therapeutics [11-14], and cancer cells that have acquired drug resistance [12, 18].

In the current study, we investigated changes in the membrane lipids and proteins of melanoma cells, while acquiring drug-resistance to dabrafenib, and when incubated with anti-melanoma peptides inspired by HDP. We selected two cyclic peptides, cyclic tachyplesin I (cTI) and cyclic gomesin (cGm), previously shown to have antimelanoma properties [19-22], to investigate whether they can kill drug-tolerant and drug-resistant melanoma cells, and if they can avoid acquired drug resistance. We explored anticancer properties of cTI and cGm against metastatic BRAF V600E melanoma cells, while they acquired resistance to dabrafenib and found that cTI and cGm display similar efficacy against melanoma cells that are drug-naïve, drug-tolerant, or drug-resistant. Melanoma cells treated with cTI over 14 weeks remained peptide-sensitive, suggesting that cancer cells do not develop resistance against these anticancer peptides. Overall, these results demonstrate that membrane-active cyclic peptides inspired by HDPs can target drug-tolerant and drug-resistant melanoma cells, are less likely to induce drug resistance than targeted therapy with kinase inhibitors, and therefore have the potential to be developed as anticancer and/or as adjuvant agents.

## Methods

### Peptide synthesis and characterisation

The synthesis, folding and purification of the peptides used in this study were done as previously described for cTI [22] and cGm [23, 20]. Briefly, both peptides were synthesized by Fmoc solid-phase chemistry on an automatic peptide synthesizer (Symphony, Protein Technologies Inc., Tucson, USA) [24]. 2-chlorotrityl resin was used for the synthesis of both cyclic peptides, which were oxidized overnight in 0.1 M ammonium bicarbonate buffer at pH 8.5 and purified using reverse-phase (RP)-HPLC (solvent A: H_2_O, 0.05% (v/v) trifluoroacetic acid (TFA), solvent B: 90% (v/v) acetonitrile, 0.05% (v/v) TFA) to a purity of >95%. The mass of the folded/reduced peptides was confirmed by ESI-MS, while their correct folding was confirmed using 1D NMR spectroscopy on a Bruker Avance 600 MHz spectrometer. Peptides purity of ≥ 95 % was confirmed by analytical RP-HPLC and their concentration was determined by absorbance at 280 nm. Estimated extinction coefficient based on the contribution of Tyr and Trp residues, and disulfide bonds was ɛ280 = 8730 M^-1^.cm^-1^ and 3230 M^-1^.cm^-1^ for cTI and cGm, respectively.

### Cell culture

WM164 and HT144 melanoma cell lines were authenticated by short tandem repeat profiling, and most recently in February 2022, and were regularly checked for absence of mycoplasma contamination using TRI mycoplasma core facility. Cells were grown in RPMI 1640 medium supplemented with 5% (v/v) fetal bovine serum (FBS; heat inactivated for 30 min at 56 °C) and with 1-2% (v/v) penicillin/streptomycin grown in tissue culture flasks in an incubator set at 37 °C with 5% CO_2_. Cells were subcultured by dilution upon reaching ∼90% confluence, every 2-3 days.

### Resistance to dabrafenib

Drug-tolerant persisters (DTPs) were generated as before [9]. In brief, WM164 and HT144 cells were grown in cell culture flasks (T175) in the presence of 100 nM dabrafenib (from 1 mM stock in DMSO) (GSK2118436; Selleckchem; a BRAF V600E-specific small molecule inhibitor). The medium containing dabrafenib was exchanged every 3-4 days. Three flasks were maintained for each cell line, and cells were harvested by trypsinisation to conduct lipid extractions, to test cytotoxicity assays, or for culture maintenance to prevent overcrowding. Cell scrapers were used to harvest cells used in proteomic studies.

### Resistance to cyclic tachyplesin (cTI)

WM164 and HT144 cells were grown in T25 flasks to ∼80% confluency in duplicates. Cells were grown in RPMI 1640 supplemented with 5% (v/v) FBS and 2% (v/v) Penicillin/Streptomycin and treated with increasing concentrations of cTI over a 14-week time course. Cells were treated with 0.5 µM of cTI added to the culture medium, increasing by an additional 0.5 µM every two weeks until 3 µM cTI, which was maintained for the last four weeks. Fresh medium containing freshly added peptide (from 1.5-3 mM stock) was replaced every 2-3 days. Cells were harvested for cytotoxicity assays and lipid extractions.

### Cytotoxicity assays

Cytotoxicity was done as previously described [22]. Briefly, cells were seeded at 5×10^3^ cells/well in a 96-well plate the day before the assay. Serum-free medium (90 µL) and peptide (10 µL) were added as two-fold dilution series (final peptide concentration ranged from 0.25 to 16 µM). Medium with 5% FBS was used when the assay was read 72 h after the addition of peptides. Blank with Dulbecco’s phosphate-buffered saline (DPBS; pH 7.0-7.3, 138 mM NaCl, 2.7 mM KCl, 8.1 mM Na_2_HPO_4_, 1.5 mM KH_2_PO_4_) and control with Triton X-100 at final concentration of 0.01% (v/v) were included to establish 0% and 100% of cell death, respectively. After 2 h incubation at 37 °C, 10 µL of sterile 0.05% (w/v) resazurin solution was added to each well and incubated overnight. The conversion of resazurin to the pink and fluorescent compound resorufin by viable cells [25] was measured in a CLARIOstar Plus microplate reader (λ_exc_ = 558 nm and λ_em_ = 587 nm).

The percentage of cell death was calculated by comparison of the fluorescence emission signal obtained with cells treated with DPBS or with Triton X-100. Dose-response curves were done with at least three biological replicates. The concentration required to inhibit the proliferation of 50% of the cells (IC_50_) was calculated by fitting dose-response curves using saturation binding with Hill slope (H): Cell death (%) = 100×[peptide]^H^/(IC_50_^H^+[peptide]^H^). Curves were fitted on Prism (Graphpad Software, Inc).

Cell membrane integrity was monitored after 40 min incubation with peptides, by quantifying the activity of lactate dehydrogenase (LDH) released into the supernatant using a colorimetric assay (CytoTox 96® Non-Radioactive Cytotoxicity Assay kit from Promega) and following the manufacturer’s instructions.

### Microscopy

Microscopy images were taken on a CKX41 inverted microscope (Olympus) with 4x, 10x, 20x magnification. The contrast and saturation of the images were enhanced using Fiji [26].

### Real-time cell analysis followed with xCELLigence

Real-time cell proliferation was monitored using an xCELLigence RTCA DP instrument (Agilent Technologies) with RTCA 16-well E-plate and placed inside an incubator at 37 °C and with 5% CO_2_. Medium was added to the plates to measure the background signal, followed by 10^4^ cells per well, which were left to settle in the plate for an hour at room temperature in the biosafety cabinet. The plates were returned to the xCELLigence and cell proliferation was monitored every 15 min for 24 h. The next day, medium was replaced with serum-free medium before peptide addition. Plates were returned to the xCELLigence and cell proliferation was monitored every 5 min for the first 3 h followed by every 30 min for 72 h.

### Cell surface charge measured using Zeta Potential

Cells grown to 80% confluency were harvested using Gibco® cell dissociation buffer, centrifuged, counted, and resuspended in cold DPBS at 2.5×10^5^ cells/mL, and stored on ice before measurement. Disposable zeta cuvettes (Malvern, Folded capillary zeta cell) were rinsed with 100% ethanol, then ddH_2_O followed by DPBS. Cuvettes were filled with cell suspension and placed in the zetasizer (Malvern, Zetasizer Nano ZS) and equilibrated for 500 s at 37 °C. Twelve measurements were acquired at a voltage of 40 V, with 50 sub runs and 90 s pause between each run. The experiments were conducted in three biological replicates using cell suspensions obtained from independent cell passages.

### Lipid extraction from cell membranes

Lipid extracts were obtained following a method based on Matyash *et al*. (2008) and adapted by Young, *et al*. (2021). Briefly, cells were trypsinized and counted using an automated TC-20 cell counter (BioRad) and triplicates of 10^6^ cells were pelleted in microfuge tubes, washed with DPBS twice, and stored at −80 °C before extractions. Lipid membrane extraction was performed in 2 mL glass vials and a stock solvent mix containing 370 µL methyl tert-butyl ether (MTBE) with 0.01% butylated hydroxytoluene (BHT) and 20 µL SPLASH LIPIDOMIX® Mass Spec Standard (Avanti Polar Lipids) per 10^6^ cells to enable quantification of lipids on mass spectrometry. Cell pellets were homogenized in 110 µL of methanol and 390 µL of solvent mix by vortexing samples for 20 s, followed by continuous agitation for 1 h. Phase separation was performed by adding 100 µL ammonium acetate (150 mM), by vortexing samples for 20 s followed by centrifugation for 5 min at 2,000 x g. The upper (organic) phase was transferred to a clean 2 mL glass vial and stored at −80 °C before analysis. The experiments were conducted in three biological replicates using cell pellets obtained from independent cell passages.

### Lipidome analysis

Tandem mass spectrometry (MS) of the lipids was performed using a triple quadrupole mass spectrometer (6500 QTRAP, SCIEX, ON, Canada) and methods were adapted from Young, *et al*. (2021). Briefly, lipid extracts were diluted 100-fold in LC-MS grade methanol prior to analysis. A 100 µL loop injection of the diluted lipid sample was entrained into a continuous flow of 5 mM methanolic ammonium acetate (NH_4_OAc) and directly infused into the electrospray ionization source of the mass spectrometer at 15 µL/min.

Using precursor ion or neutral loss scans, tandem mass spectrometry was performed for classed-based lipid species detection (PC and SM: precursor ion scan *m/z* 184.2, collision energy [CE] = 39 V; PE: neutral loss scan *m/z* 141.1, CE = 29 V; PS: neutral loss scan *m/z* 185.2, CE = 29 V; PI neutral loss scan *m/z* 277, CE = 32 V). Transfer capillary temperature was held constant at 150 °C throughout, with curtain and collision (CAD) gasses set to 20 (arb) and 10 (arb), respectively. Delustering, entry and exit potentials were set to 50 V, 11 V and 16 V, respectively. Lipidview® (Version 1.3 beta; SCIEX) software was used for data processing of Sciex data files obtained from the QTRAP 6500, including referencing biological lipids to the lipid class-based, deuterated internal standards and correction of abundances to account for naturally occurring isotopic lipids. Statistical analysis, including graphical visualisation of the data, was conducted using Microsoft Excel.

### Cholesterol quantification by derivatization method

Free cholesterol from cell lipid extracts was quantified using a method adapted from Liebisch *et al*. (2006). Briefly, cell lipid extracts (50 µL) were dried under nitrogen (N_2_) gas before adding 500 µL of chloroform and 100 µL of acetyl chloride to convert free cholesterol to cholesteryl acetate (CE 2:0). Samples were mixed, left at room temperature for 1 h and dried under N_2_. Dried lipid samples were redissolved in 1 mL methanolic NH_4_OAc (10 mM) and stored at −20 °C. 50 µL samples were loop injected in the electrospray ionization source of the mass spectrometer (6500 QTRAP, SCIEX, ON, Canada) using a flow of methanolic NH_4_OAc (5 mM). Quantification was achieved by combining selected reaction monitoring (SRM) transitions of *m/z* 446.4 > 369.4 (CE 2:0) for cholesterol and 453.4 > 376.4 (D_7_-CE 2:0) for the deuterated D_7_-cholesterol internal standard.

### Whole cell lysate and protein digestion for proteomics

Cells were lysed with lysing buffer containing 1% sodium deoxycholate in 100 mM Tris pH 8.5, 10 mM tris(2-carboxyethyl) phosphine and 40 mM 2-chloroacetamide. Samples were sonicated for 20 min in a pre-cooled Bioruptor (Diagenode) and centrifuged at 13,000 x g for 10 min to remove insoluble particles. Total protein concentration was determined using Direct Detect Infrared Spectrometer (Merck) and a solution with 20 µg of protein was heated at 95 °C for 5 min. Protein samples were digested overnight with 400 ng of trypsin at 37 °C. Reactions were stopped using 10% (v/v) trifluoroacetic acid (TFA) (0.5% (v/v) final concentration) and samples were centrifuged for 10 min at 13,000 x g to remove precipitates. Cleaved peptides were cleaned using C18 tips and eluted in 80% (v/v) acetonitrile. Samples were vacuum dried and resuspended in 0.5% (v/v) TFA before LC-MS/MS analysis.

### Proteome analysis

Digested proteins were resolved on an online U3000 RSLCnano nanoHPLC system with a 50 cm Easy-Spray C18 analytical column (Thermo Fisher, catalogue 160454 and ES803A) and analysed on a Q Exactive™ Plus Orbitrap™ Mass spectrometer (Thermo Scientific, USA). Mobile phases were solvent A: 0.1% (v/v) formic acid (FA) in H_2_O, and solvent B: 80% (v/v) acetonitrile with 0.1% (v/v) FA in H_2_O. Peptide samples of 1 µg were loaded in 3% of solvent B, and eluted using a gradient of 3% to 30% of solvent B during 144 min, followed by a gradient of 30% to 50% of solvent B over 20 min at a flow rate of 250 nL/min. Peptides were analysed using positive ionization mode using settings typical for high complexity peptide analyses. Raw data analysis was performed using Proteome Discoverer (version 2.4.1.15) for protein identification searching against the *Homo sapiens* proteome (SwissProt Taxonomy 9606, database release v2020 10 18) and an in-house assembled protein-contaminant database.

### Peptide-membrane binding affinity monitored using surface plasmon resonance

Surface plasmon resonance (SPR) was used to investigate binding of cTI and cGm to lipid bilayers composed of lipid membrane extracts obtained from WM164 cells in drug-naïve, DTPs (treated with dabrafenib for 30 days) and permanent drug resistant (PDR) state (treated with dabrafenib for 105 days). Lipid membrane extracts were obtained as above described, and organic solvents were dried under nitrogen gas flow and vacuum desiccation overnight to obtain a lipid film. A suspension with lipid vesicles was obtained by hydrating the lipid film with SPR running buffer (10 mM HEPES, 150 mM NaCl, pH 7.4, filtered with 0.20 µm pore size membrane), and vortexing. The vesicles were sized to obtain small unilamellar vesicles (SUVs, diameter of 50 nm) using freeze/thaw and extrusion through membranes with 50 nm pores as previously detailed [28].

Measurements were conducted using a Bioacore T200, sensor L1 chips and with controlled temperature at 25 °C. SUVs were deposited onto the L1 chip at a flow rate of 2 µL/min for 60 min to achieve coverage of the chip surface with a lipid bilayer, as confirmed with a plateau in the signal. Serial dilutions of peptide samples prepared in running buffer were injected over the deposited lipid bilayers at 5 µL/min for 180 s and the dissociation was monitored for 600 s. Washing and regeneration of the chip were done as before [28]. Data normalisation, dose response curves, determination of peptide-to-lipid ratio when peptide-lipid binding reaches saturation (P/L_max_), and determination of the membrane partition coefficient (K_p_), (i.e., the ratio of molar fractions of peptide molecules in the lipid and aqueous phases when peptide-lipid association/dissociation reaches equilibrium) were calculated for each peptide-membrane extract as previously detailed and validated [29].

## Results

### Induction of drug resistance to dabrafenib in melanoma cell lines

To generate drug-resistant melanoma cells, we incubated BRAF V600E positive metastatic melanoma cell lines WM164 and HT144 with 100 nM dabrafenib until they reached permanent drug resistance (≥ 90 days) (Fig 1a&b). Both melanoma cell lines were monitored by microscopy during development of acquired drug resistance (Fig 1b). Treatment of both cell lines with dabrafenib caused death of drug-naïve cells within 48 h. A population of slow-cycling cells, referred to as DTPs [3, 9], survived (Fig. 1a&b I) and re-entered active growth from 30 days onwards, which can be visualized by the formation of colonies (Fig. 1a&b, II). After the colony state, cells were able to continuously proliferate in the presence of dabrafenib (Fig. 1a&b, III), suggesting that both cell lines became permanently resistant. The emergence of permanent drug-resistant cells (PDR cells, PDRCs) was monitored in three biological replicates for each cell line.

**Fig. 1.**
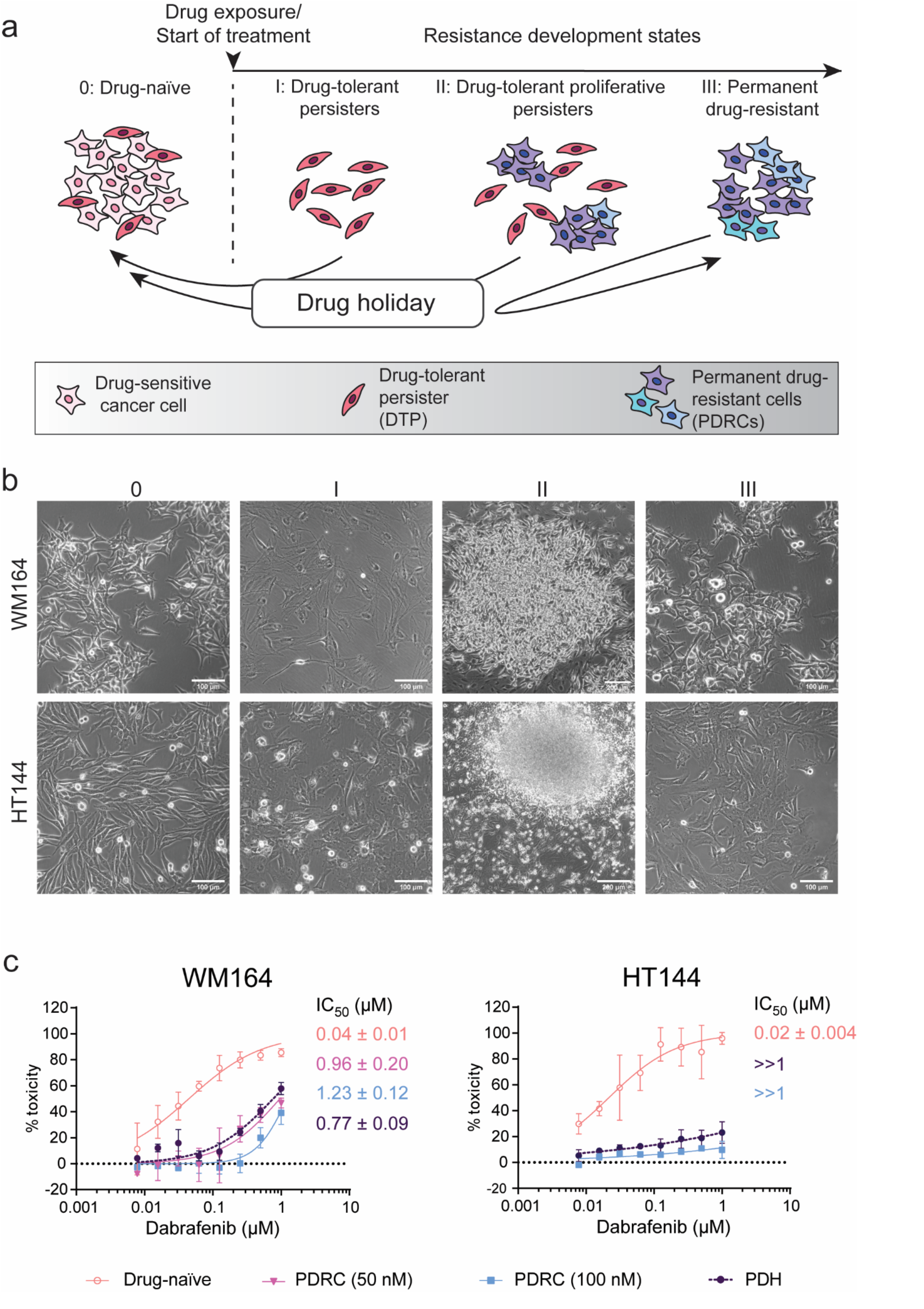
Development of melanoma cells with acquired resistance to dabrafenib. **a**. Diagram illustrating development of adaptive resistance to targeted therapy; Drug-naïve: in a cancer cell population dynamic phenotypic switching occurs between proliferative drug sensitive and slow-cycling cells; Drug-tolerant persisters (DTP): After commencement of treatment, slow-cycling drug-tolerant cells undergo epigenetic reprogramming. When drug is removed for two weeks (drug holiday), these cells return to a drug-naïve state; Drug-tolerant proliferative persisters (DTPPs): Drug-tolerant subpopulations re-enter proliferation; Permanent drug resistant (PDR): Emergence of permanent drug-resistant cells (PDRCs) that will remain resistant after a drug holiday. Adapted from [3]. **b**. Microscopy images of WM164 (top row) and HT144 (bottom row) cells during treatment with 100 nM dabrafenib at the drug-naïve (0), DTP (I), DTPP (II) and PDR (III) state. Scale bars represent 100 µm for panel 0, I, III and 200 µm for panel II. **c**. Dabrafenib cytotoxicity toward drug-naïve and dabrafenib-resistant WM164 and HT144 melanoma cells. PDRCs were incubated for more than 100 days with 100 nM dabrafenib (PDRC (100 nM)) or with 50 nM dabrafenib (PDRC (50 nM)). After 100 days with 100 nM dabrafenib, cells were allowed a “drug holiday” for two weeks and were tested post-drug holiday (PDH). Resazurin assays were performed three days after addition of drug and dose response curves were obtained from three independent replicates each with cells generated from three independent experiments. Data points represent mean ± SD. Using one-way ANOVA, IC_50_ of WM164 (both with 50 nM and 100 nM dabrafenib treatment) and HT144 PDRCs compared to respective PDH cells were non-statistically significant

We confirmed that WM164 and HT144 cells became permanently resistant to dabrafenib by testing their susceptibility to this small-molecule inhibitor (Fig. 1c). Parental WM164 cells were more susceptible to dabrafenib, than PDR WM164 cells, as quantified by an IC_50_ (peptide concentration required to kill 50% of cells) of 44 nM and 1.23 µM, respectively. Moreover, PDR WM164 cells generated from treatment with 100 nM compared to previously generated PDRCs from treatment with 50 nM dabrafenib [30] had similar IC_50_ (IC_50_ of 1.23 µM and 0.96 µM, respectively). Parental HT144 cells were also more susceptible to dabrafenib (IC_50_ = 22 nM), than PDR HT144 cells (IC_50_ >> 1 µM with ∼10% toxicity at 1µM).

Permanent resistance to dabrafenib was further confirmed by testing tolerance to the drug after growing the PDRCs for two weeks without drug (i.e., drug holiday). Toxicity studies show that susceptibility of post-drug holiday cells to dabrafenib (PDH, IC_50_ = 0.77 µM; Fig. 1c) was not significantly different from that of PDRCs, which validated their permanent resistance to dabrafenib.

### Melanoma membrane lipid composition changes during the acquisition of resistance to dabrafenib

Cancer progression and development of drug resistance to anticancer treatments involve changes in lipid composition and metabolism [31-33]. In the context of this study, we are interested in investigating changes in structural lipids (i.e., sterols, sphingolipids and glycerophospholipids) that make up the lipid bilayer in cell membranes. The proportion of each structural lipid, the respective glycerophospholipid headgroups, and the length (i.e., number of carbons) and degree of unsaturation (i.e., the number of carbon-carbon double bonds) in their fatty acid (FA) chains, impacts the overall biophysical properties of the cell membrane, such as charge, fluidity, presence of raft domains and lipid bilayer thickness. Therefore, variations in the lipid composition can modulate cell membrane permeation as drug-resistance mechanisms [34, 33], and can also affect the activity of anticancer peptides targeting cancer cell membranes.

We investigated the lipid membrane composition of the tested melanoma cells while acquiring resistance to dabrafenib. To capture all states of resistance development (i.e., drug-naïve, DTPs, late persisters (DTPPs) and PDRCs) we collected WM164 and HT144 cells at different time points (0, 7 days, and every 15 days between 15 and 105 days) of treatment with dabrafenib. We extracted their lipids and quantified the proportion of the four major classes of glycerophospholipids (GPL) (i.e., phospholipids containing phosphatidylcholine (PC), phosphatidylethanolamine (PE), phosphatidylinositol (PI), or phosphatidylserine (PS)-headgroups) and of the sphingolipid sphingomyelin (SM). Further, we assigned the total number of carbons and degree of unsaturation (i.e., lipid sum composition) using mass spectrometry-based shotgun lipidomic methodologies previously described [35]. We also quantified the cholesterol content from the same lipid extracts via a mass spectrometry method adapted from Liebisch, *et al*. (2006).

The lipid composition of untreated WM164 and HT144 cells (Fig. 2a, 0 d) shows a large proportion of PC (57 and 59 mol%, respectively), followed by PE (21 and 15 mol%), PI (12% for both), PS (7 and 9 mol%) and SM (3 and 5 mol%). Incubation with dabrafenib resulted in changes in the proportion of these lipids in the persister state (Fig. 2a, 30 d); after incubation with dabrafenib for 30 days, WM164 cells show a 3.7-fold increase in SM (3 to 11 mol%), 1.2-fold increase in negatively charged lipids (PS+PI) (19 to 23 mol%), and a 0.8-fold decrease in the proportion of the zwitterionic PE and PC (78 to 66 mol%). Incubation of HT144 cells with dabrafenib for 30 days (Fig. 2a, 30d) resulted in a 1.8-fold and 1.3-fold increase in SM (5 to 9 mol%) and PE (15 to 20 mol%), respectively, whereas PC proportion decreased by 0.9-fold (59 to 51 mol%). The overall GPL and SM composition of PDR WM164 and HT144 cells is similar to respective drug-naïve cells (see 0 d and 105 d in Fig. 2a).

**Fig. 2.**
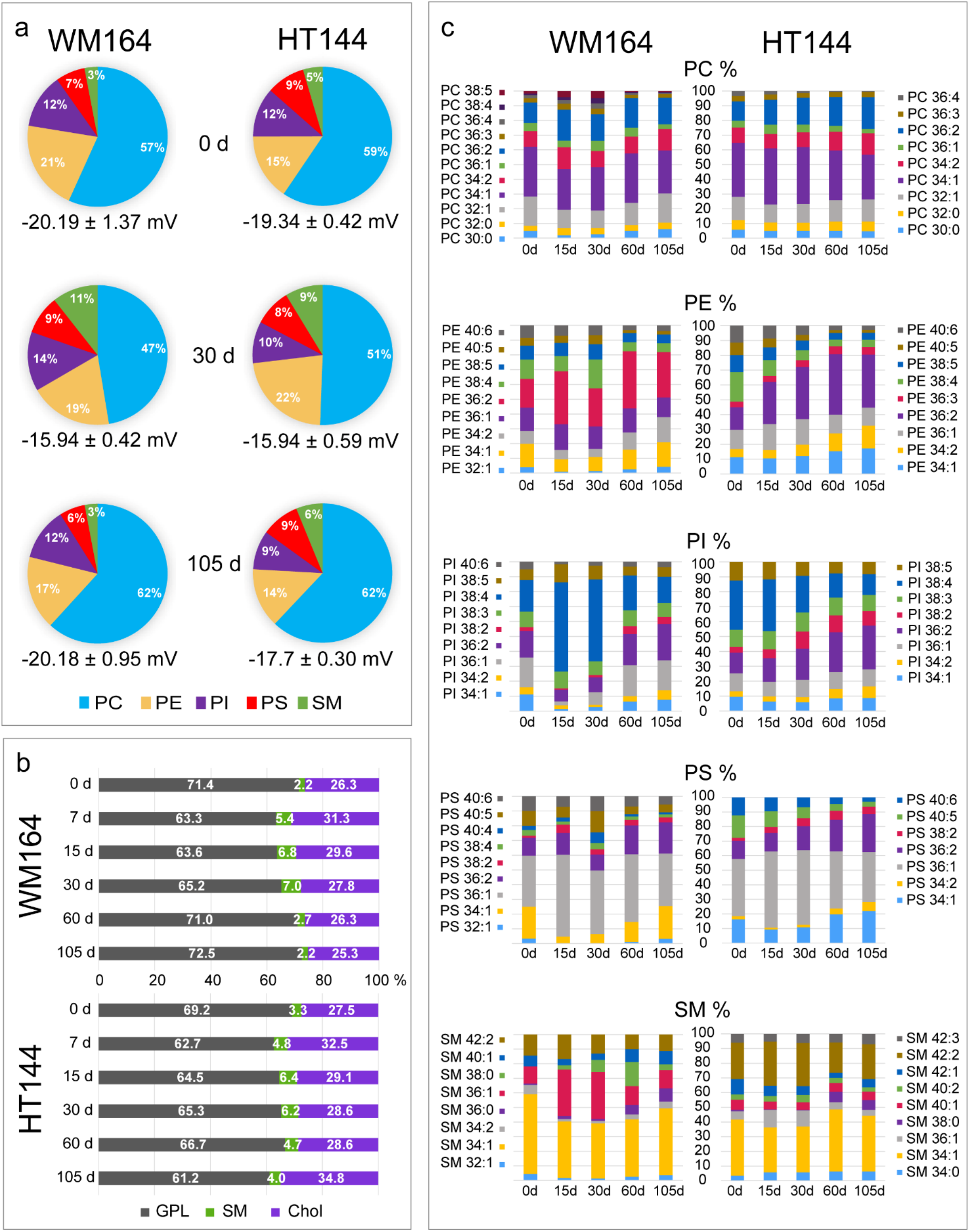
Lipid composition of WM164 and HT144 cells while acquiring resistance to dabrafenib. **a**. Pie charts show the mole percentage (mol%) of the four major glycerophospholipid (GPL) classes in mammalian cell membranes (i.e., phosphatidylcholine (PC), phosphatidyl ethanolamine (PE), phosphatidylinositol (PI) and phosphatidylserine (PS)), and sphingomyelin (SM) in WM164 and HT144 cells during treatment with dabrafenib (0, 30 and 105 days). Average surface charge (mV ± SEM) of cells treated with dabrafenib for 0, 30 and 105 days, as measured using zeta potential (250,000 cells/sample, (n=3)). See also Fig. S1. **b**. Histograms represent the normalized abundance of PC, PE, PI, PS, and SM species quantified at the sum composition level. See also file SF1. **c**. Mole percentage (mol%) of glycerophospholipids (GPL = PC + PE + PS + PI), sphingomyelin (SM) and cholesterol (Chol) during incubation with dabrafenib. Mean ratio displayed (n = 3). See also Table S1.

The proportion of cholesterol in WM164 cells was higher in DTP, but similar in PDR cells, when compared to drug-naïve cells, (see 0, 7 and 105 d of treatment with dabrafenib in Fig. 2b & Table S1). In HT144 cells, the proportion of cholesterol was higher in DTP cells and in PDR cells (see 0, 7 and 105 d in Fig. 2b & Table S1).

Comparison of the FAs in each GPL class and SM during treatment with dabrafenib, revealed changes in the degree of saturation and acyl chain length up to 30 days (Fig. 2c, see 0, 15 and 30 d, file SF1). In WM164, the proportion of shorter FA chains species in all classes decreased while longer and more unsaturated FA chains increased. For example, after 15-30 days of treatment with dabrafenib, SM 36:1 (8 to 26 mol%) and PI 38:4 (18 to 55 mol%) increased 3-fold, whereas PI 36:1, 36:2, 34:1 and 34:2 decreased. Overall, these results suggest elongation and desaturation of GPL and SM while WM164 cells are in the DTP state. WM164 cells in PDR state regained a FA profile similar to drug-naïve cells.

In HT144 cells the FA chains followed a gradual variation over time, and differences are more evident in the PDRCs, when compared to the drug-naïve cells (Fig. 2c, file SF1). For instance, long polyunsaturated (> 2 double bonds) species within the PE, PI, PS classes decreased proportionately, whereas the shorter-chain species increased over time; specifically, PE 38:4, PI 38:4, PS 40:5 were reduced by 50 mol%, whereas the proportion of PE 34:2, PE 36:2, PI 36:2 and PS 36:2 increased 2-3 fold. Trends in PC and SM were either more subtle or non-existent.

In general, DTPs display increased fractions of SM and cholesterol in both cell lines suggesting that the cell membrane of DTPs is more rigid [37, 38] than in drug-naïve cells. Untreated and PDR WM164 cells have similar lipid composition, whereas PDR HT144 cells display a decrease of polyunsaturated FA chains, and an increase in the total amount of cholesterol in comparison with untreated HT144 suggesting that the cell membrane of HT144 cells is more rigid [39] when drug-resistant.

### Surface charge of drug-treated melanoma cells

We investigated the cell surface charge of drug naïve, DTP and PDR WM164 and HT144 cells by measuring their zeta potential (i.e., the electric potential at the cell interface) when treated with dabrafenib (Fig. 2a). Cancer cells are characterised by a negative electric charge at the cell surface due to a larger proportion of negatively charged GPL (i.e. PS) exposed at the cell surface, compared to healthy cells [40, 41], which contains mainly zwitterionic and neutral lipids (i.e., PC, SM and Chol) in the outer layer [42]. However, it is not known if the surface charge is maintained when cancer cells become DTP or PDR.

Both drug-naïve cell types had negatively charged cell surfaces (WM164: −20.19 ± 1.37 mV, HT144: −19.34 ± 0.42 mV), which became less negative in DTPs (WM164: −15.94 ± 0.42 mV, HT144: −15.94 ± 0.59 mV). However, negative surface charge was regained by PDRCs (WM164: −20.18 ± 0.95 mV, HT144: −17.7 ± 0.30 mV).

The zeta-potential results suggest that DTPs have a lower amount of negatively charged lipids on the cell surface, than drug-naïve and drug-resistant cells. Comparison of the overall proportion of PI and PS in WM164 cells shows that drug-naïve (0 d, 19%) and drug-resistant cells (105 d, 18%) have similar fraction of negatively charged phospholipids, but lower amounts than DTPs (30 d, 23%). This was similar for HT144 cells (Fig. 2a), where drug-naïve cells (0 d, 21%) had a higher proportion of negatively charged lipids compared to DTPs (30 d, 18%) and PDRCs (105 d, 18%). Thus, compared to drug-naïve cells, the less-negative surface charge in the DTP state for both WM164 and HT144 cells does not correlate with a decreased fraction of anionic lipids. Overall, these results suggest that DTPs may regain an asymmetric distribution of lipids across the cell membrane leaflets, with a lower proportion of negatively charged lipids in the outer leaflet and/or contribution of other charged molecules at the cell surface (e.g., glycosaminoglycans).

### Changes in the proteome of melanoma cells in DTP and PDR states and correlation with respective lipidome

We investigated changes in the proteome while WM164 and HT144 cells developed resistance towards dabrafenib. We collected total protein extracts and performed whole cell proteomic studies on drug-naïve, DTP and PDR WM164 and HT144 cells. Cells treated with dabrafenib were compared with drug-naïve cells and processed following the same protocol (i.e., duration of cell culture, cell harvesting, whole-cell lysate, protein digestion, protein recovery and amount of total protein).

When compared to drug-naïve cells, DTPs had more proteins that were differentially expressed (Fig. 3a) than PDRCs in both cell lines. Using Kyoto Encyclopedia of Genes and Genomes (KEGG) pathway (Fig. S2) [43-45] and gene ontology (GO) (Fig. S3), we identified overexpression of proteins that have been proposed to be involved in drug resistance mechanisms, such as lysosomal proteins and degradation of fatty acids (see more details in SI and in Fig. S3). Relevant for this study, we also identified overexpression of proteins in pathways that are likely to modulate cell membrane properties; specifically, proteins involved in the degradation of glycosaminoglycans, and in lipid metabolism pathways (i.e. biosynthesis of glycosphingolipids and PI, and degradation of sphingolipids), as detailed below.

**Fig. 3.**
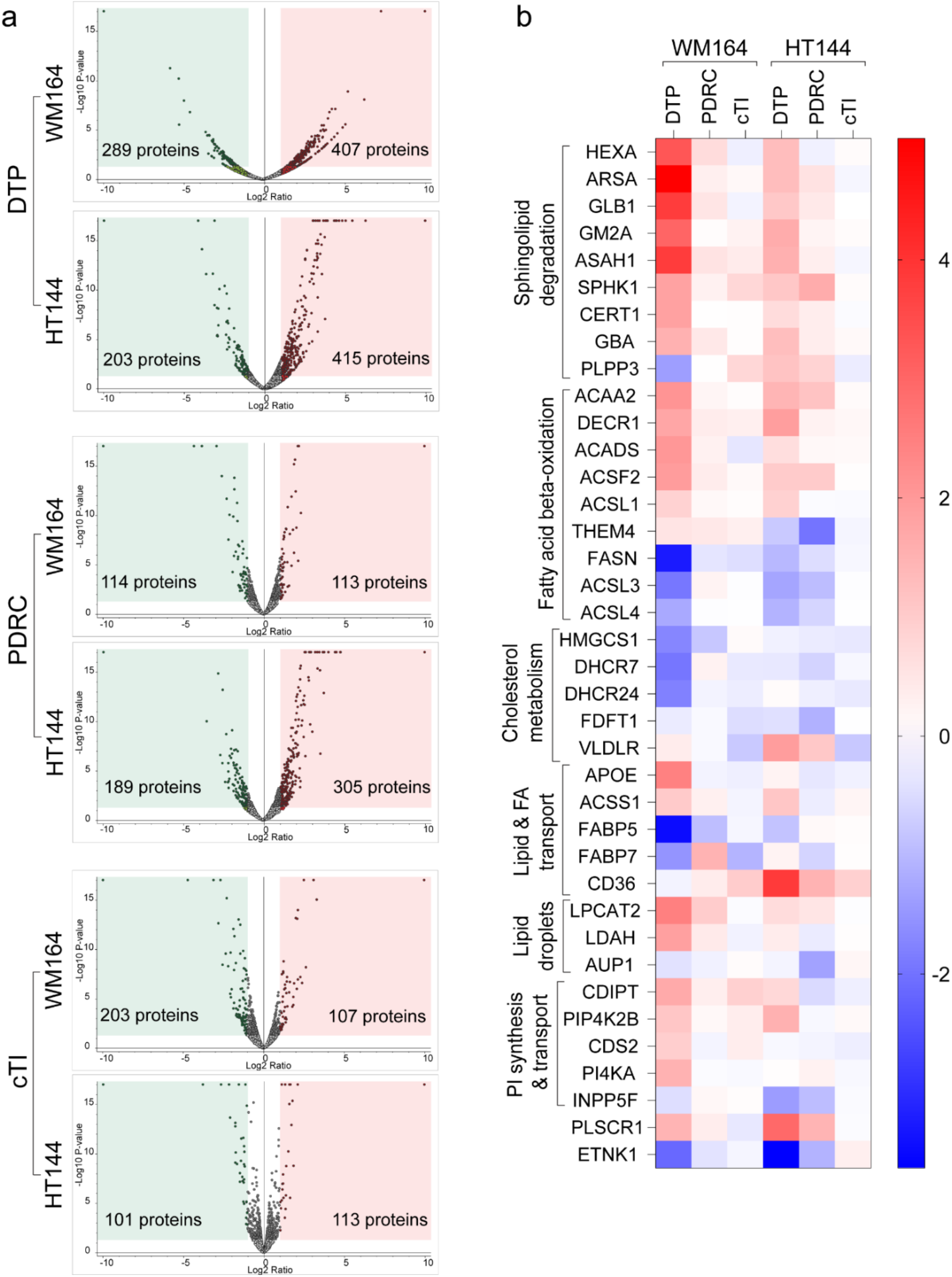
Proteomic data of WM164 and HT144 cells treated with cTI. **a**. Volcano plots of significantly downregulated (green) and upregulated (red) proteins of WM164 and HT144 cells in a drug-tolerant state (DTP; 28 days with 100 nM dabrafenib), drug-resistant state (PDRC; ≥ 105 days with 100 nM dabrafenib), and of cells treated with cTI for 13 weeks (cTI), compared to untreated cells. *p*-value ≤ 0.05; log2 fold change ± 1. See also Fig. S2 and S3. **b**. Heat map of downregulated (blue) and upregulated (red) proteins in significant lipid synthesis and metabolic pathways. See also file SF2.

Degradation of glycosaminoglycans and other glycans is supported by the enrichment in hydrolase activity on glycosyl bonds/compounds in WM164 DTPs (Fig. S3). These proteins and polysaccharides have been shown to play critical roles in cancer proliferation, invasion and metastasis, and in therapeutic resistance [46, 47]. Glycosaminoglycans are negatively charged and thus their degradation might contribute to the less-negative surface charge observed in the DTPs’ zeta potential.

Proteins involved in the biosynthesis of glycosphingolipids (KEGG: hsa00603 & hsa00604) and in the degradation of sphingolipids (e.g., HEXA, GLB1, ARSA, ASAH1 and GBA [48] are upregulated in DTPs, but not in PDRCs (Fig. 3b, Fig. S2). Sphingolipid degradation leads to increased ceramide abundance, which can be converted into SM in a reaction catalysed by sphingomyelin synthase [49]. Changes in the proteins involved in sphingolipid degradation are likely to contribute to the increased proportion of SM detected in the lipidome of DTPs.

Proteins that play a role in the PI biosynthesis, such as CDIPT, CDS2, PI4KA and PIP4K2B, are upregulated in the DTP state, especially in WM164 cells (Fig. 3b, file SF2). Upregulation of PI biosynthesis can explain the relatively high amount of phosphatidylinositol (PI 38:4) detected in the lipidome. PI 38:4 is preferentially converted to phosphatidylinositol polyphosphates such as phosphatidylinositol 3,4,5-trisphosphate (PIP3) [50]. PIP3 activates protein kinase B (AKT), a downstream effecter of PI3 kinase signalling, which regulates multiple pathways involved in multidrug resistance, including inhibition of apoptosis, cell growth and metabolism stimulation [51].

Most proteins associated with cholesterol biosynthetic pathways were downregulated (e.g. HMGCS1, DHCR7, DHCR24, FDFT1) in WM164 DTPs, whereas alipoprotein E (APOE), which is involved in cholesterol transport and lipid metabolism [52, 53], was overexpressed by 5.6-fold (log2: 2.48) (Fig. 3b, file SF2). In PDR WM164 cells, the expression level of proteins involved in cholesterol biosynthetic pathways is similar to that of drug-naïve cells (fold change between 0.6 to 1.2). In HT144 cells, the expression of those proteins was in general lower in both DTPs and PDRCs (fold change between 0.5 and 0.9), except for the very low-density lipoprotein receptor (VLDLR) expression (Fig. 3b, file SF2). The larger proportion of cholesterol detected in WM164 and in HT144 DTPs might be obtained from increased uptake of cholesterol from the medium (mediated by APOE in WM164, and by VLDLR in HT144 [54]) as opposed to being synthesised by the cells.

### cTI and cGm, membrane-active peptides with anti-melanoma properties

As the lipid membrane composition of WM164 and HT144 cells changed while cells developed resistance to dabrafenib, we wanted to examine if membrane-active peptides could maintain anticancer activity towards melanoma cells with acquired drug-resistance. The positively charged HDPs tachyplesin I (TI) from the horseshoe crab *Tachypleus tridentatus* and gomesin (Gm) from the spider *Acanthoscurria gomesiana* and their respective ultra-stable cyclic analogues, cyclic tachyplesin (cTI) and cyclic gomesin (cGm) (Fig. 4a), selectively kill melanoma cells [55, 20-22]. Their mechanism of action is distinct from small molecule anticancer drugs and involves preferential binding and disruption of melanoma cell membranes, compared to non-cancerous cell membranes [19-22]. We selected the backbone cyclic peptides, cTI and cGm, which have improved drug-like properties, compared to the parent peptides, to investigate whether they could kill drug-tolerant and drug-resistant melanoma cells, and if they could avoid acquired drug resistance.

**Fig. 4.**
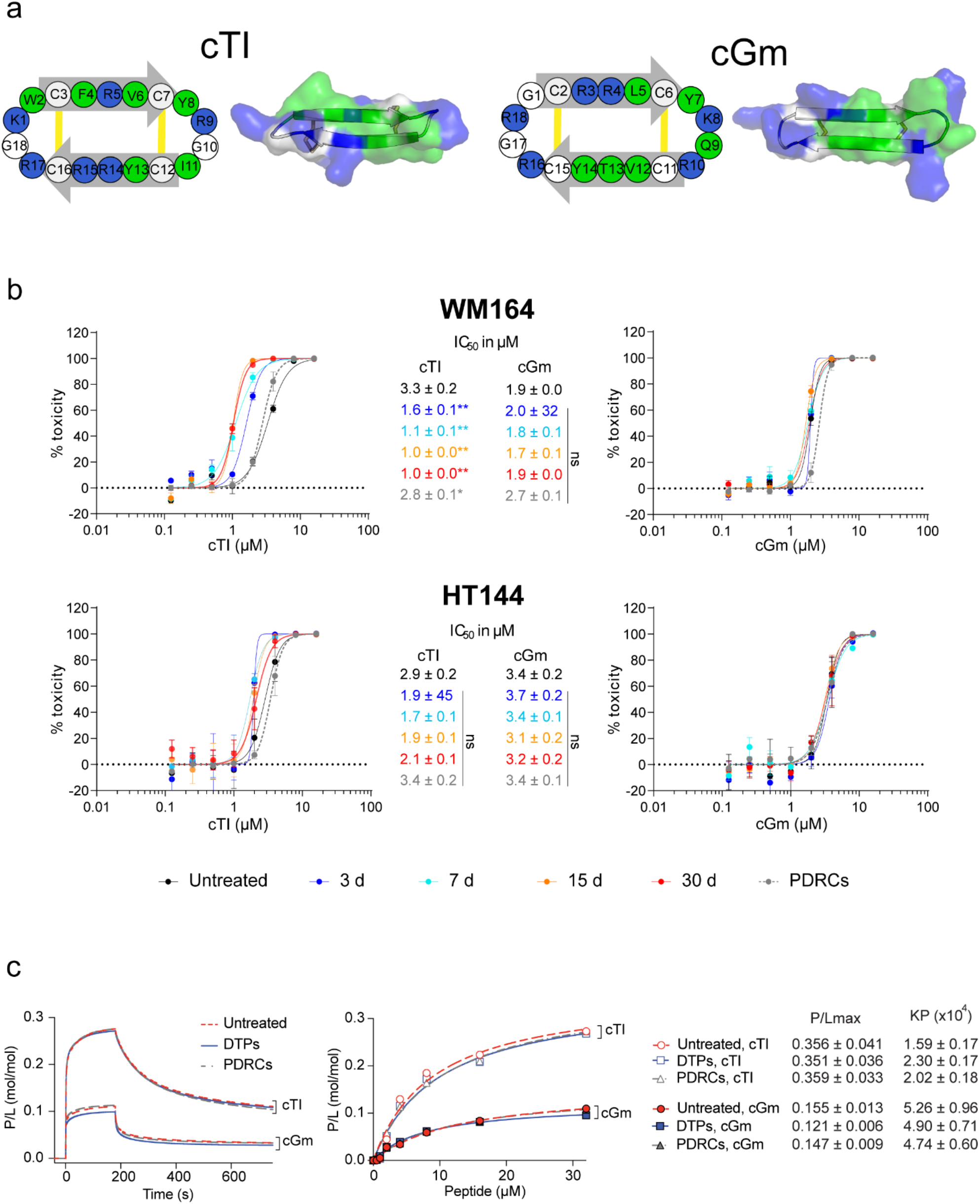
Anti-melanoma properties of cyclic beta-hairpin peptides. **a**. Schematic of the amino acid sequence (left) and three-dimensional structure (right) showing the distribution of positively charged (blue) and hydrophobic (green) amino acids of cyclic tachyplesin I (cTI) and cyclic gomesin (cGm), with an overall net charge of +7 and +6, respectively. Both peptides possess a β-hairpin structure with two antiparallel β-strands (grey arrows) stabilized by two disulfide bonds (yellow). **b**. Cytotoxicity of cTI and cGm toward WM164 and HT144 cells while acquiring resistance to dabrafenib: cells in drug-naïve (untreated), drug-tolerant persisters (DTP; 3, 7, 15 and 30 d), or with acquired permanent drug resistant (PDRCs; 105 d) states. Peptides were incubated with cells for 24 h, and toxicity was quantified by measuring conversion of resazurin into fluorescent resorufin. Dose response curves were obtained from three independent experiments. Data points represent mean ± SEM. Significance determined by one-way ANOVA by comparing fitted IC_50_ ± SEM of untreated with treated cells: not significant (ns) p > 0.05, * p ≤ 0.01, and ** p ≤ 0.0001. **c**. Binding of cGm and cTI to lipid bilayers prepared with lipid membrane extracts obtained from WM164 untreated cells, DTPs (30 d) and PDRCs (105 d) monitored using surface plasmon resonance. Peptide samples with a range of concentrations (0-32 µM) were injected over lipid bilayers deposited onto L1 chip for 180 s (association phase), and dissociation was monitored for 600 s (dissociation phase). Left panel shows representative sensorgrams obtained with peptide at 32 µM. Response units were converted to peptide-lipid ratio (P/L (mol/mol)). Right panel shows peptide-lipid binding dose-response curves: P/L obtained at the end of association phase (i.e., when the binding phase reaches a steady state and using t =170 s as a reporting point) plotted in function of peptide concentration. Curves were fitted in GraphPad Prism 8 using saturation binding with Hill slope. P/L_max_ is the peptide-to-lipid ratio when binding reaches saturation and was determined from the fitting P/L dose response curves. Membrane partition coefficient, Kp, is the ratio of molar fractions of peptide in the peptide and aqueous phase and was determined by fitting dose-response data using a partition formalism [56].

The toxicity of cTI and cGm was tested against drug-naïve (0 d), DTP (i.e., after 3, 7, 15 and 30 d of incubation with 100 nM dabrafenib), and PDR (after 105 d of incubation with 100 nM dabrafenib) WM164 and HT144 cells (Fig. 4b). Dose-response curves show that cGm and cTI displayed toxicity towards drug-naïve, DTP and PDRCs with the IC_50_ range of 1.0-3.3 µM against WM164 cells and of 1.7-3.7 µM against HT144 cells. cGm has similar efficacy for killing cells in different resistance states whereas the efficacy of cTI varies; nevertheless, the cytotoxicity data for WM164 and HT144 cells demonstrate the ability of cTI and cGm to kill drug-naïve, drug-tolerant and drug-resistant melanoma cells.

### cTI and cGm peptides kill DTPs & PDRCs via a membrane disruption mechanism

We studied the binding affinity of cTI and cGm to lipid bilayers prepared with lipids extracted from drug-naïve, DTP or PDR WM164 cells using surface plasmon resonance. Sensorgrams, dose-response curves and fitted parameters (Fig. 4c) show that cTI had similar high binding affinity for bilayers composed of lipids extracted from WM164 cells before, during and after acquiring drug resistance to dabrafenib, as quantified by the fitted peptide-lipid binding at saturation (P/L_max_), and membrane partition coefficient (K_P_). A similar binding affinity of cGm for lipid membranes obtained from drug-naïve, DTP or PDR WM164 cell membrane extracts was observed (Fig. 4c). Comparison between the two peptides show that cTI reached a higher P/L_max_ and dissociated slower from bilayers composed of lipids extracted from WM164 membranes, than cGm. These results show that cTI has a higher affinity for those membranes than cGm. The high peptide-lipid binding affinity obtained with cell membrane extracts agrees with previous results with bilayers formed with synthetic lipids showing high affinity for lipid membranes containing negatively-charged phospholipids [19-21]. and demonstrates that both cTI and cGm bind membranes composed of lipids extracted from drug-naïve, drug-tolerant and drug-resistant melanoma cells.

To investigate the ability of these peptides to eradicate drug-naïve and drug-resistant melanoma cells via a mechanism involving cell membrane disruption, we monitored the proliferation of WM164 and of HT144 cells in real time by recording their electrical impedance (Fig. S4) using the xCELLigence real time cell analysis. Treatment of cells with 4 µM cGm or with 5 µM cTI for 24 h, resulted in ∼50% reduction in impedance signal, and the highest concentration of peptide tested (i.e., 8 µM cGm, or 10 µM cTI) displayed a drop in the impedance to ∼0 within 2-3 hours, indicating rapid cell death; the dose effect and rapid cell death suggest a mechanism involving cell membrane disruption [57]. Similar trends were observed with cells that were drug-naïve or drug-resistant to dabrafenib. These results agree with the toxicity studies and the respective IC_50_ obtained considering number of cells used in each assay (e.g., in xCELLigence 10^4^ WM164 cells seeded for 24 h and treated with 5 µM cTI resulted in ∼ 50% of cell proliferation, whereas in cytotoxicity studies using resazurin 5 x10^3^ WM164 cells seeded for 24 h and treated with 3 µM of cTI resulted in 50% of cell death).

To confirm that both cGm and cTI induce cell membrane disruption in WM164 DTPs, we followed the enzymatic activity of cytosolic lactate dehydrogenase (LDH) released into the media. The LDH activity was measured after incubation with either peptide for 40 min and compared to a maximum achieved by treating cells with Triton X-100. The concentration of peptide required to release 50% of the maximum LDH activity (cTI LDH_50_: 1.5 ± 0.2 µM, cGm LDH_50_: 2.3 ± 0.2 µM; data not shown) was near identical to the inhibitory concentrations (see Fig. 4; cTI IC_50_: 1.3 ± 0.2 µM, cGm IC_50_: 2.1 ± 0.2 µM). This confirmed that both cTI and cGm eradicated DTPs and PDRCs by a rapid mechanism that involved cell membrane disruption.

To determine whether co-treatment of dabrafenib with either peptides had an additive or synergetic effect, drug-naïve and PDR WM164 cells were co-treated with dabrafenib and cTI or cGm (see further details in SI and data in Fig. S5). The results show that the combination of peptide with drug had an additive effect.

### Melanoma cell lines do not develop resistance following continued exposure to cTI

We selected cTI as a candidate to investigate whether melanoma cells develop tolerance, or resistance, to peptides as it has a higher binding affinity toward melanoma cell membrane. We exposed WM164 and HT144 melanoma cell lines to increasing concentrations of cTI over time, starting from 0.5 µM and increasing an additional 0.5 µM every 15 days until total concentration was 3 µM (WM164 IC_50_ ∼3.2 µM; HT144 IC_50_ ∼2.5 µM). Cells were incubated with 3 µM for another 30 days. Changes in cell morphology were monitored via microscopy (Fig. 5a) and the susceptibility of cells to cTI was monitored by quantifying toxicity using a resazurin assay (Fig. 5b).

**Fig. 5.**
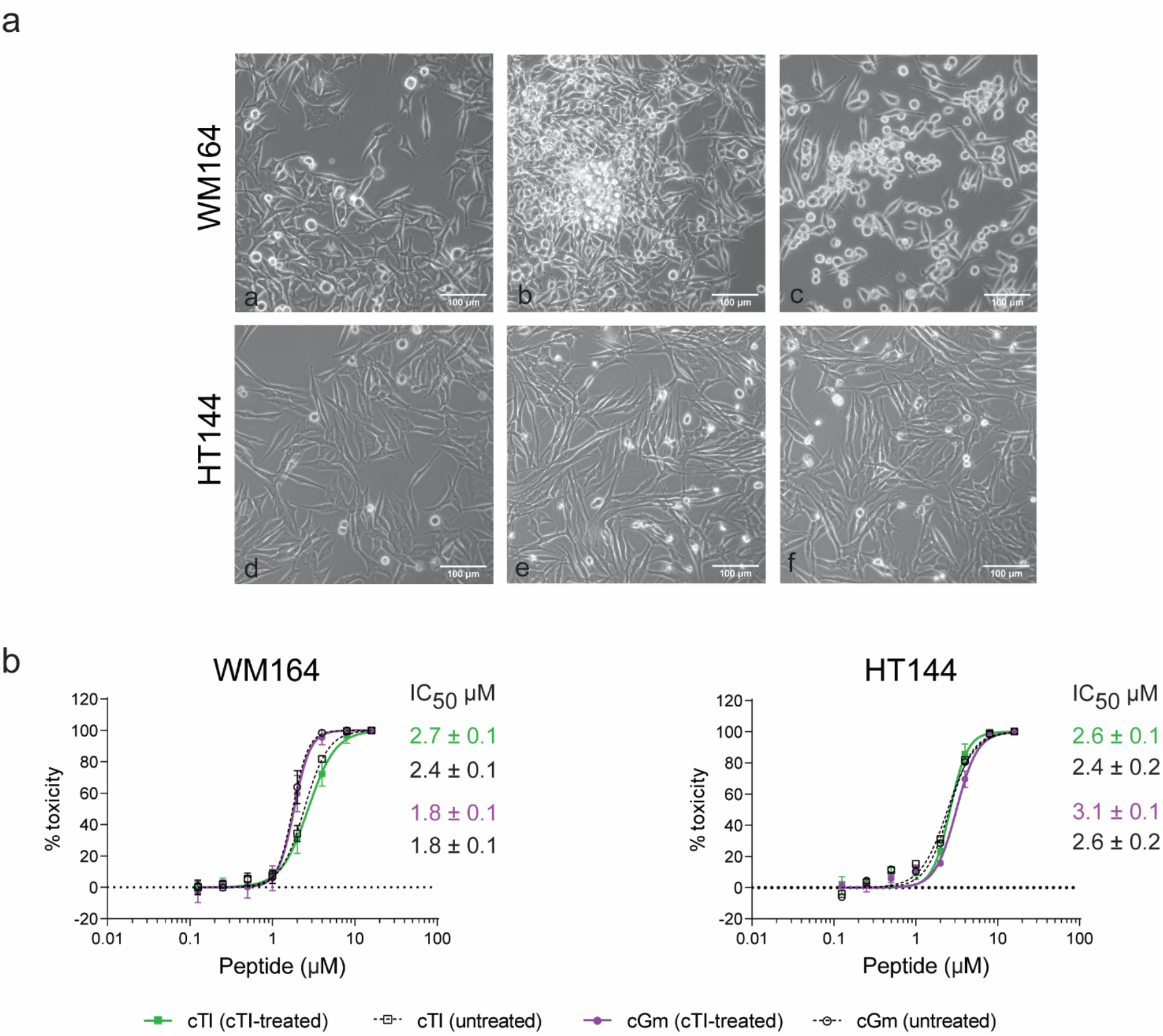
Treatment of melanoma cells with cTI for 100 days. **a**. Micrographs obtained with WM164 and HT144 cells at day 0 without peptide (**a**,**d**), at 50 days with 2 µM cTI (**b**,**e**), and at 100 days with 3 µM cTI in growth medium (**c**,**f**). **b**. Dose-response curves of cGm and cTI on cells treated with cTI for 13 weeks. WM164 and HT144 cells were incubated with peptide for 24 h and cell death was measured using resazurin assay. Data points represent mean ± SEM of two (HT144) or three (WM164) biological replicates, except untreated cell control where data points are from one (HT144) or two (WM164) replicates which did not change over 100 days

The morphology of WM164 cells did not change until the concentration of cTI reached 2 µM (50 days). Some cell clusters, like the colonies observed with the incubation with dabrafenib (phase II), were visible; however, the cells in these clusters were rounder in appearance. After incubation with 2 µM cTI for one week, WM164 cells became more elongated and less dendritic, and more dead cells were detected floating in the media. After continuously treating cells with cTI for 100 days, the cells were grown without the peptide for 3 weeks to mimic a “drug holiday”. After two days without cTI, the morphology of WM164 cells was similar to that of the untreated cells. In contrast to observations for WM164 cells, the morphology of HT144 cells remained unchanged throughout treatment with cTI for 100 days.

Cytotoxic assays were performed throughout the continuous treatment of WM164 and HT144 cells with cTI to monitor whether cells maintained susceptibility to cTI. Near identical dose-response curves as quantified by Hill slope and IC_50_ obtained with untreated cells and cells treated with cTI for 100 days supports a similar efficacy and mechanism (Fig. 5b). Untreated and cTI-treated cells were equally susceptible to cGm, as confirmed by IC_50_ and dose-response curve. Overall, these studies confirm that WM164 and HT144 cells did not develop resistance towards cTI and remain susceptible to cTI and a similar peptide, even after treatment for extended period.

### Overall lipid composition of melanoma cell membranes unaffected by cTI treatment

Using a similar approach to the lipidomic analysis of cells while acquiring resistance to dabrafenib, we characterised the lipidome of cTI-treated melanoma cells during the 100-day treatment regimen (Fig. 6a). Comparison with untreated cells displayed minimal differences in the proportion of the different lipid classes detected with the exception of PE. WM164 cells treated with 2 µM cTI had an increase in the proportion of PE from 15 in the untreated cells to 23 mol% after incubation for 100 days. In contrast, HT144 treated with 2 µM cTI had an increased proportion from 16 in the untreated cells to 20 mol% at 60 days and a subsequent reduction to 16 mol% after 100 days of treatment. The increase in the proportion of PE in the membrane might explain the morphological changes detected, as PE-phospholipids are known to have a role in inducing membrane curvature [58].

**Fig. 6.**
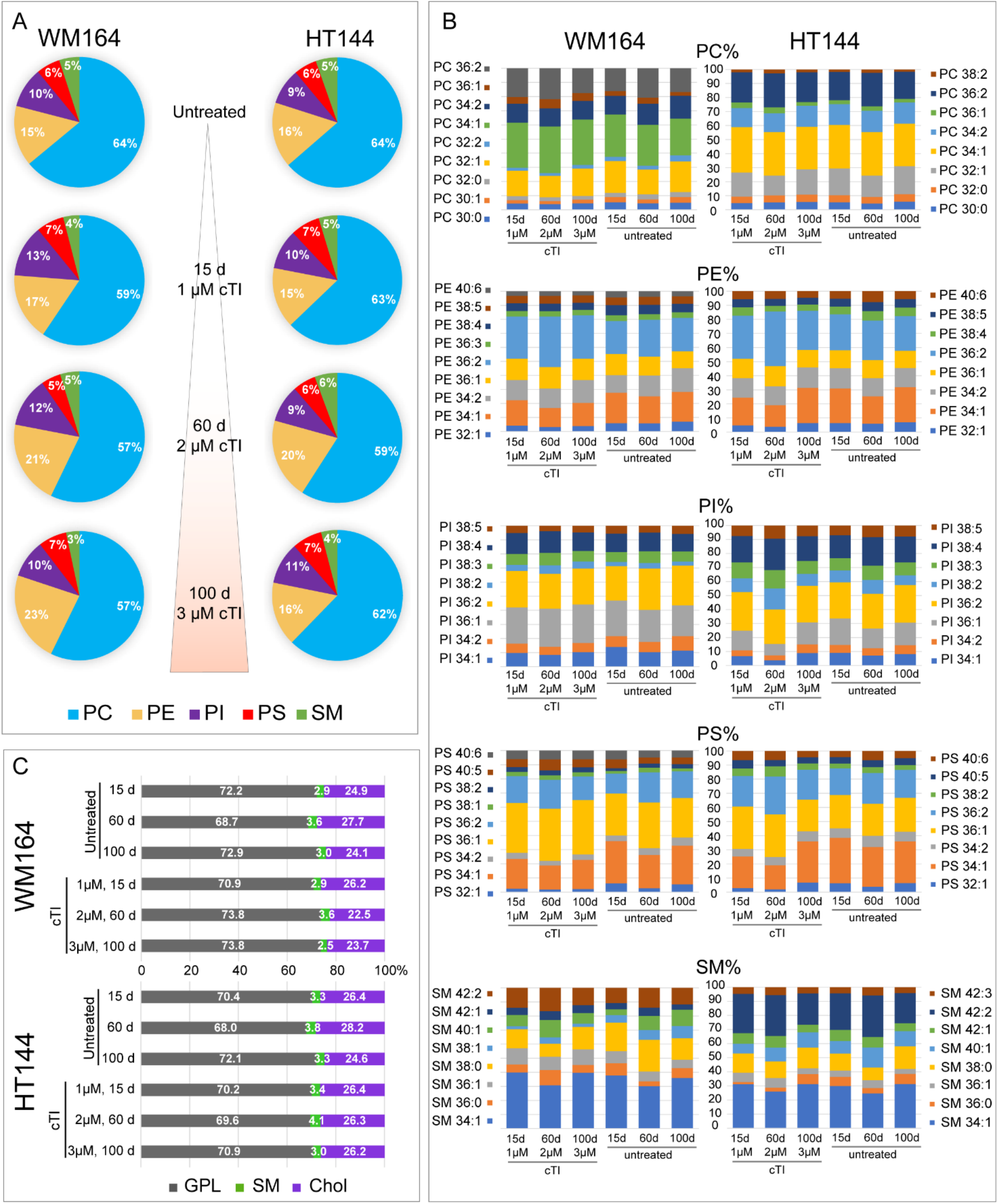
Proportion of glycerophospholipid classes and sphingomyelin, and respective fatty acid chains in WM164 and HT144 during exposure to increasing concentrations of cTI over a period of 14 weeks. **a**. Pie charts represent the proportion (mol%) of PC, PE, PI, PS, SM of WM164 and HT144 cells untreated and treated with 1 µM (15 days), 2 µM (60 days) and 3 µM (100 days) cTI. See also Fig. S6. **b**. Histograms represent the normalized abundance of PC, PE, PI, PS, and SM species quantified at the sum composition level. See also file SF3. **c**. Mole percentage (mol%) of glycerophospholipids (GPL = PC, PE, PS, PI), sphingomyelin (SM) and cholesterol (Chol) during incubation with cTI. Mean ratio displayed (n = 2). See also Table S2

WM164 cells treated with cTI over time displayed only slight variations in the acyl chain composition of each lipid class, when compared with control untreated cells (Fig. 6b). The lipid sum composition of untreated HT144 cells differed from those incubated with cTI for 60 days, while at 100 days untreated cells and those treated with cTI had very similar lipid sum compositions across all monitored lipid classes. Variations in the proportion of cholesterol in untreated and cTI-treated cells over time did not follow a particular trend for either WM164 or HT144 cells (Fig. 6c & Table S2).

Although cTI eradicates melanoma cells by targeting and disrupting the lipid bilayer in the cell membrane, the tested melanoma cells do not change their membrane composition in the presence of cTI, in contrast to the observation for cells treated with the BRAF inhibitor dabrafenib. This result further supports the toxicity studies and indicates that melanoma cells do not develop resistance towards cTI.

### cTI did not affect expression of proteins involved with membrane lipid composition

We performed proteomic studies on WM164 and HT144 cells that were incubated for 100 days with cTI to examine whether any changes occurred in the proteome compared to untreated melanoma cells.

Differences in the expression of proteins involved in lipid metabolism were not evident in melanoma cells treated with cTI; the proteome data (Fig. 3) supported the lipidomics results (Fig. 6), which suggest that cTI does not induce changes in expression of structural lipids that compose cell membranes. Nevertheless, GO functional analysis of proteins from cTI-treated melanoma cells showed that changes on the cell periphery and membrane were accompanied by downregulation in proteins involved in the ECM structure (Fig. S3). These results support the action of cTI on the cell membrane, consistent with previous observations, but also suggest that cTI does not impact protein expression for pathways that could promote resistance to this peptide.

## Discussion

The development of resistance to anticancer drugs is a recurring problem during cancer treatment, and this is particularly true for patients with metastatic melanoma that carry a BRAF V600E mutation. It has been proposed that slow-cycling cells are responsible for the ability of a tumor to survive and adapt to drug exposure [8, 59, 9, 60]. Thus, drugs able to kill slow-cycling cells are of utmost importance for preventing cancer progression. Here we induced tolerance to dabrafenib, a small molecule drug that targets tumor cells with BRAF V600E mutation, in metastatic melanoma cell lines WM164 and HT144 to produce slow-cycling and drug-resistant cells, and investigated their lipid composition, the role of lipid metabolism in acquired drug-resistance and their susceptibility to membrane-active peptides cTI and cGm.

Alterations in the lipid metabolism and of membrane lipid composition can contribute to acquired drug-resistance from treatment with targeted and conventional therapies [31]. Here, we confirmed that treatment with dabrafenib induces metabolic reprogramming, as demonstrated by alterations in the lipid composition and differential expression of proteins involved in lipid biosynthesis and FAO.

It is not clear whether these changes occurred to protect cancer cells from stress induced by dabrafenib, or were inherited from a pre-existing subpopulation of drug-resistant cancer cells; however, these changes were not observed following sustained treatment with cTI.

Lipid composition results demonstrate increased proportion of SM and cholesterol, and/or a decrease in long polyunsaturated lipids in HT144 DTPs; all these changes are likely to increase membrane rigidity [61, 62, 33] and reduce cellular uptake of dabrafenib. Changes in membrane fluidity has been reported as a drug-resistance mechanism in similar studies with lung [63], breast [34, 64], and colorectal cancers [65].

Alterations in the membrane of slow-cycling melanoma cells could also affect the activity of membrane-active peptides, such as cTI and cGm; however, here we demonstrated that both peptides rapidly kill drug naïve, slow-cycling and drug-resistant melanoma cells through cell membrane disruption (see Fig. 4b and Fig. S4). A mechanism dependent on membrane agrees with previous studies showing disruption of negatively charged cell membranes [19-22]. Differences in their membrane interaction and/or preference for specific membrane components might explain the higher efficacy of cTI, compared to cGm, and against DTPs, compared to drug-naïve WM164 cells. Although there are some modifications in the total lipid composition of DTPs and PDRCs, we do not know how the lipids are distributed within the lipid bilayer of cell membranes and across organelle membranes, nor how variations in other membrane components exposed at the cell surface (e.g., glusosaminoglycans) might contribute to the potency of these peptides.

We monitored the effect of continuous exposure to cTI on the susceptibility of WM164 and HT144 cells to cTI or cGm peptides. The treated cells were equally susceptible to these peptides compared to untreated cells, suggesting that melanoma cells did not acquire resistance to cTI treatment over 100 days. The current study, together with other studies combining treatment of TI peptides with other anticancer drugs (e.g., cisplatin) [66], and the absence of observed drug-related adverse effects in mouse [67] and rat models [68, 69] with tachyplesin III, support the notion that stable cTI- and cGm-analogues could find application as additional treatment to small-molecule kinase inhibitors to eradicate the slow-cycling cancer cell population during anticancer treatment, before resistance to the kinase inhibitor can develop, and without inducing resistance themselves.

### Significance

Acquired drug-resistance is a serious problem for the treatment of patients with metastatic melanoma; melanoma tumors contain slow-cycling cells that are tolerant to treatment with small molecule drugs targeting the RAF/MEK pathway and are very difficult to eradicate, which lead to acquired drug-resistance. Here, we demonstrate the application of cGm and cTI, stable membrane-active peptides with anti-melanoma properties, as alternative antimelanoma agents. These cyclic peptides kill slow-cycling and drug-resistant melanoma cells via a fast mechanism, and do not induce resistance. Therefore, cyclic peptides are excellent candidates for developing new anticancer therapies with the potential to target slow-cycling cancer cell subpopulations thereby preventing the emergence of resistance mechanisms.

## Acknowledgements

Peptides used in this study were synthesised using facilities from the ARC Centre of Excellence for Innovations in Peptide and Protein Science managed by David Craik group (IMB, UQ, Brisbane, Australia), lipidomics data were acquired at the Central Analytical Research Facility (CARF) at the Queensland University of Technology (Brisbane, Australia) and the authors acknowledge technical support from Berwyck Poad and Pawel Sadowski (CARF, QUT). Proteomics data were obtained at the Proteomics Facility at the Translational Research Institute (TRI, Brisbane, Australia). The authors thank Olivier Cheneval and Joachim Weidmann (IMB, UQ, Brisbane, Australia) for help with the synthesis of the initial batch of peptides.

## Author contributions

A.B., F.V., H.S., N.L., S.J.B., S.T.H. contributed to the study design and conception. A.B., F.V., R.S.E.Y., S.J.B., N.L., H.H., H.S., and S.T.H. developed methodology. A.B, F.V., R.S.E.Y, F.N.B., S.T.H., did the experimental work, analyzed, and interpreted data. A.B., F.V. and S.T.H. wrote the manuscript, with input from N.L., H.S., R.S.E.Y, S.J.B. and D.J.C. All the authors read the manuscript and contributed specific expertise. S.T.H., D.J.C., and S.J.B. acquired funding and provided resources to conduct the experimental work of this study.

## Consent to publish

All authors have read and approved the final version of the manuscript.

## Data availability

All data generated and analysed in this study are included in this published article and its supplementary information and files. Proteomic datasets for this study are available from the corresponding author on reasonable request.

## Supplementary files

This manuscript includes supplementary files:

- Supplementary information, including Fig. S1-S6, Tables S1 & S2.
- Supplementary file 1 (SF1): spreadsheet with quantification of lipid species – dabrafenib treatment
- Supplementary file 2 (SF2): spreadsheet with raw data of manually identified proteins
- Supplementary file 3 (SF3): spreadsheet with quantification of lipid species – cTI treatment

