## supplementary information for "Membrane-active peptides escape drug-resistance in cancer"

a

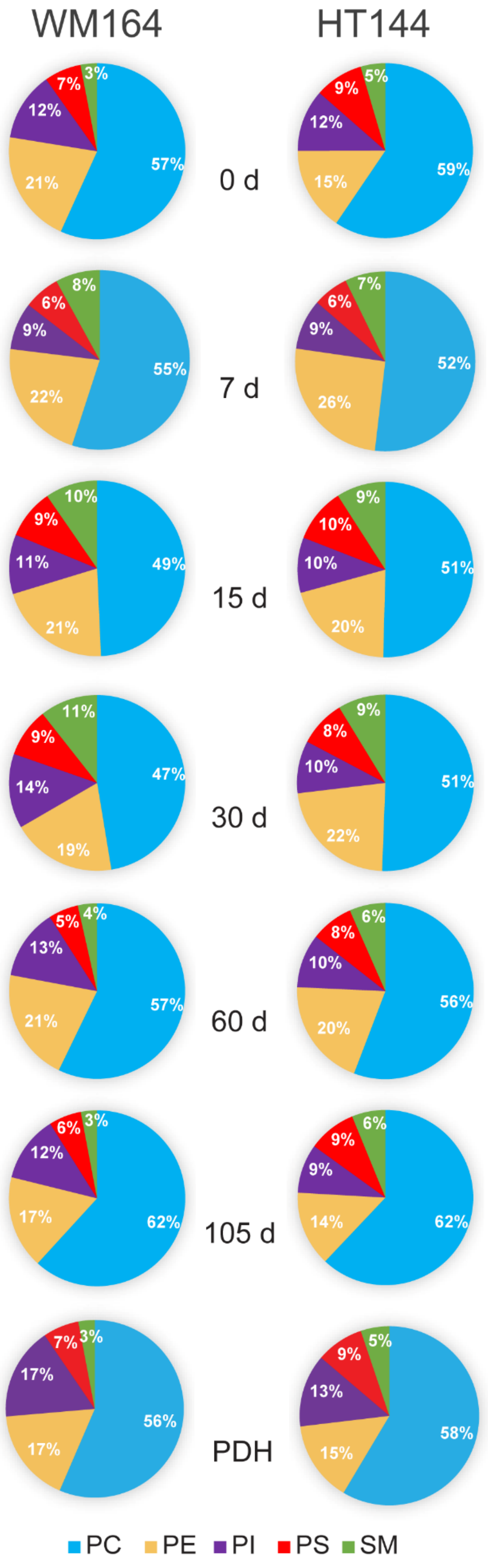


b

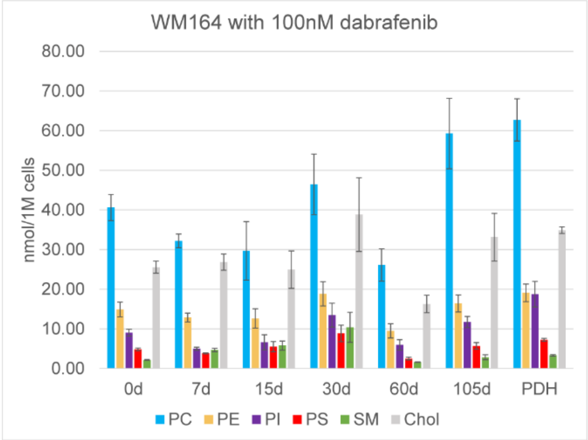


| | | 0d | 7d | 15d | 30d | 60d | 105d | PDH |
|---|---|---|---|---|---|---|---|---|
| nmol/1M cells | PC | 40.60 | 32.20 | 29.68 | 46.45 | 26.13 | 59.28 | 62.74 |
| | PE | 14.89 | 12.85 | 12.62 | 18.83 | 9.49 | 16.40 | 19.08 |
| | PI | 9.04 | 5.00 | 6.60 | 13.47 | 5.96 | 11.73 | 18.75 |
| | PS | 4.84 | 3.82 | 5.51 | 8.85 | 2.51 | 5.70 | 7.25 |
| | SM | 2.17 | 4.67 | 5.78 | 10.40 | 1.61 | 2.85 | 3.30 |
| | Chol | 25.56 | 26.85 | 24.93 | 38.84 | 16.28 | 33.12 | 34.89 |
| SEM | PC | 3.28 | 1.72 | 7.39 | 7.64 | 4.07 | 8.89 | 5.32 |
| | PE | 1.86 | 1.10 | 2.44 | 3.06 | 1.80 | 2.14 | 2.21 |
| | PI | 0.86 | 0.39 | 1.90 | 3.02 | 1.33 | 1.41 | 3.21 |
| | PS | 0.34 | 0.13 | 1.29 | 2.11 | 0.35 | 0.82 | 0.36 |
| | SM | 0.10 | 0.42 | 1.14 | 3.79 | 0.07 | 0.61 | 0.19 |
| | Chol | 1.52 | 2.04 | 4.73 | 9.31 | 2.20 | 6.03 | 0.82 |

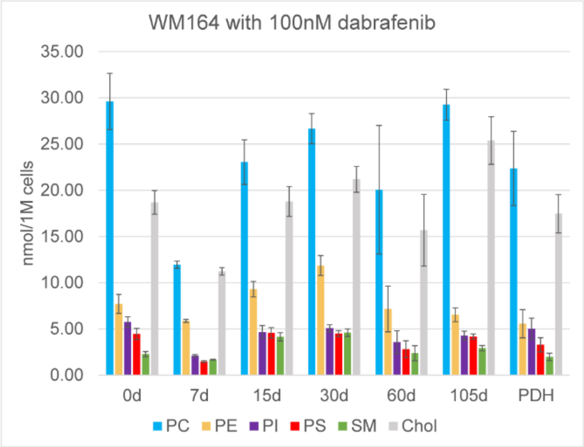


| | | 0d | 7d | 15d | 30d | 60d | 105d | PDH |
|---|---|---|---|---|---|---|---|---|
| nmol/1M cells | PC | 29.61 | 11.94 | 23.04 | 26.65 | 20.05 | 29.25 | 22.35 |
| | PE | 7.70 | 5.86 | 9.30 | 11.86 | 7.16 | 6.54 | 5.56 |
| | PI | 5.74 | 2.09 | 4.66 | 5.07 | 3.55 | 4.28 | 5.02 |
| | PS | 4.46 | 1.47 | 4.57 | 4.50 | 2.81 | 4.17 | 3.28 |
| | SM | 2.27 | 1.67 | 4.15 | 4.59 | 2.37 | 2.91 | 1.97 |
| | Chol | 18.69 | 11.23 | 18.78 | 21.17 | 15.67 | 25.38 | 17.46 |
| SEM | PC | 3.04 | 0.38 | 2.41 | 1.63 | 6.95 | 1.67 | 4.01 |
| | PE | 1.03 | 0.16 | 0.84 | 1.08 | 2.47 | 0.75 | 1.53 |
| | PI | 0.57 | 0.12 | 0.70 | 0.38 | 1.25 | 0.48 | 1.15 |
| | PS | 0.60 | 0.10 | 0.57 | 0.32 | 0.91 | 0.28 | 0.77 |
| | SM | 0.28 | 0.05 | 0.44 | 0.41 | 0.82 | 0.29 | 0.41 |
| | Chol | 1.27 | 0.41 | 1.61 | 1.40 | 3.87 | 2.58 | 2.07 |

**Fig. S1. Lipid composition of WM164 and HT144 cells while acquiring resistance to dabrafenib. Related to Fig. 2. a.** Pie charts show proportion (mol%) of the four major glycerophospholipid classes in mammalian cell membrane (i.e., PC-, PE-, PI-, and PS) and of sphingomyelin (SM) in WM164 and HT144 cells during treatment with dabrafenib for 0, 7, 15, 30, 60, 105 days and 15 days without drug (PDH). **b.** Absolute quantification of each lipid class in nmol per 1 million cells. Data represented as mean ± SEM (n = 3)

#### Lipidomic profile changes

It has been suggested that a high PE:SM ratio can act as an ‘ON’ switch for proliferation signals at the plasma membrane by facilitating high Ras-MAPK activity and therefore proliferation, whereas a low PE:SM ratio leads to a denser surface membrane packing and impedes the transduction of proliferative signals [1]. WM164 cells have a lower PE:SM ratio in the DTP state (e.g., 1.7:1 at 30 d), than in the drug-naïve (7:1) or PDR (5.7:1) state; however, in HT144 cells the ratio of these lipids displayed only a small variation (e.g., 3:1 for drug naïve, 2.4:1 for DTPs at 30 d and 2.3:1 for PDRCs). The results from these two melanoma cell lines suggest that the proportion of SM in relation to all phospholipids, rather than the PE:SM ratio, is a better measure to distinguish proliferative cells (i.e., drug-naïve and PDR) from slow-cycling cells.

**Table S1:** Mole ratio of glycerophospholipids (GPL), sphingomyelin (SM) and cholesterol (Chol) during incubation of WM164 and HT144 cells with 100 nM dabrafenib over 105 days and after 15 days without dabrafenib (PDH). GPL include PC, PE, PS and PI. Mean ratio ± SEM displayed (n = 3). Statistical difference (*P*) between cholesterol ratios determined by 2-way ANOVA were non-significant for most compared conditions, except for the ones indicated with the same letters corresponding to a p-value of <0.05 (a,f), ≤ 0.01(c), ≤ 0.001(b,d), ≤ 0.0001(e). Related to Fig. 2c

| | Days with Dab | % GPL | % SM | % Chol | *P* | SEM GPL | SEM SM | SEM Chol |
|---|---|---|---|---|---|---|---|---|
| WM164 | 0 | 71.4 | 2.2 | 26.3 | | 0.7 | 0.1 | 0.6 |
| | 7 | 63.3 | 5.4 | 31.3 | ab | 0.7 | 0.2 | 0.6 |
| | 15 | 63.6 | 6.8 | 29.6 | | 1.0 | 0.2 | 0.9 |
| | 30 | 65.2 | 7.0 | 27.8 | | 2.8 | 1.7 | 1.2 |
| | 45 | 69.2 | 4.6 | 26.2 | | 1.8 | 1.3 | 0.5 |
| | 60 | 71.0 | 2.7 | 26.3 | | 1.8 | 0.3 | 1.7 |
| | 75 | 70.8 | 2.8 | 26.4 | | 1.9 | 0.5 | 1.5 |
| | 90 | 70.8 | 2.8 | 26.4 | | 2.1 | 0.3 | 1.8 |
| | 105 | 72.5 | 2.2 | 25.3 | a | 0.8 | 0.2 | 0.7 |
| | PDH (2 weeks) | 74.1 | 2.4 | 23.5 | b | 1.3 | 0.1 | 1.2 |
| HT144 | 0 | 69.2 | 3.3 | 27.5 | cdef | 0.6 | 0.1 | 0.7 |
| | 7 | 62.7 | 4.8 | 32.5 | c | 0.9 | 0.2 | 1.0 |
| | 15 | 64.5 | 6.4 | 29.1 | | 0.4 | 0.1 | 0.4 |
| | 30 | 65.3 | 6.2 | 28.6 | | 0.3 | 0.3 | 0.1 |
| | 45 | 66.4 | 5.8 | 27.8 | | 0.5 | 0.2 | 0.3 |
| | 60 | 66.7 | 4.7 | 28.6 | | 0.4 | 0.2 | 0.2 |
| | 75 | 61.7 | 5.0 | 33.3 | d | 0.9 | 0.4 | 1.3 |
| | 90 | 59.3 | 4.9 | 35.8 | e | 0.7 | 0.1 | 0.5 |
| | 105 | 61.2 | 4.0 | 34.8 | e | 1.9 | 0.1 | 1.9 |
| | PDH (2 weeks) | 64.5 | 3.5 | 32.0 | f | 2.2 | 0.2 | 0.0 |

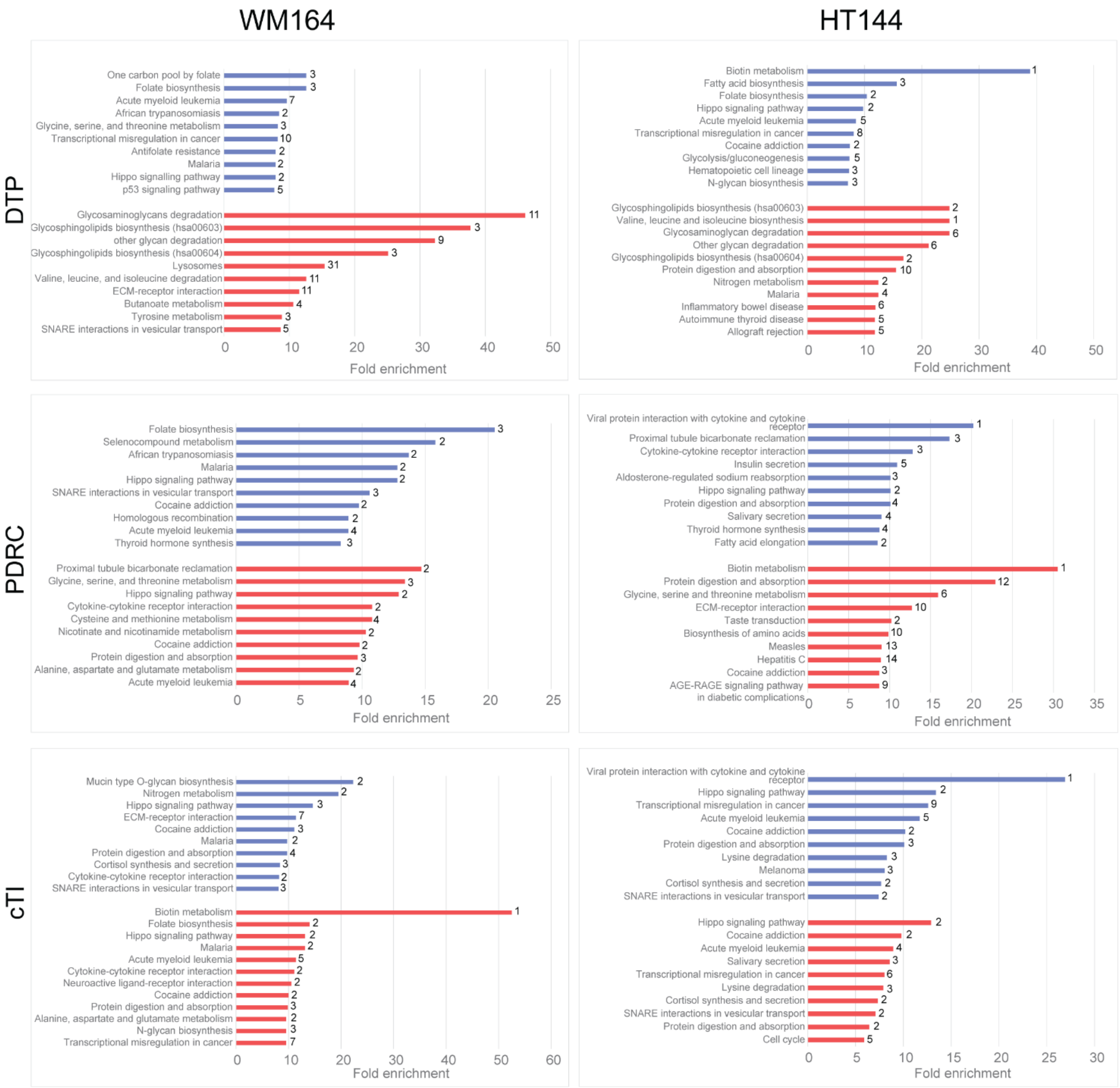


**Fig. S2 Top 10 KEGG pathways with highest fold enrichment. Related to Fig. 3.** Diagrams represent number (end of each bar) of upregulated (red) and downregulated (blue) proteins involved in identified KEGG pathways (https://www.genome.jp/kegg/pathway.html) with *p*-value ≤ 0.05; log2 fold change ± 1. (fold enrichment: enrichment factor calculated as a quotient of number of found protein and number of expected proteins)

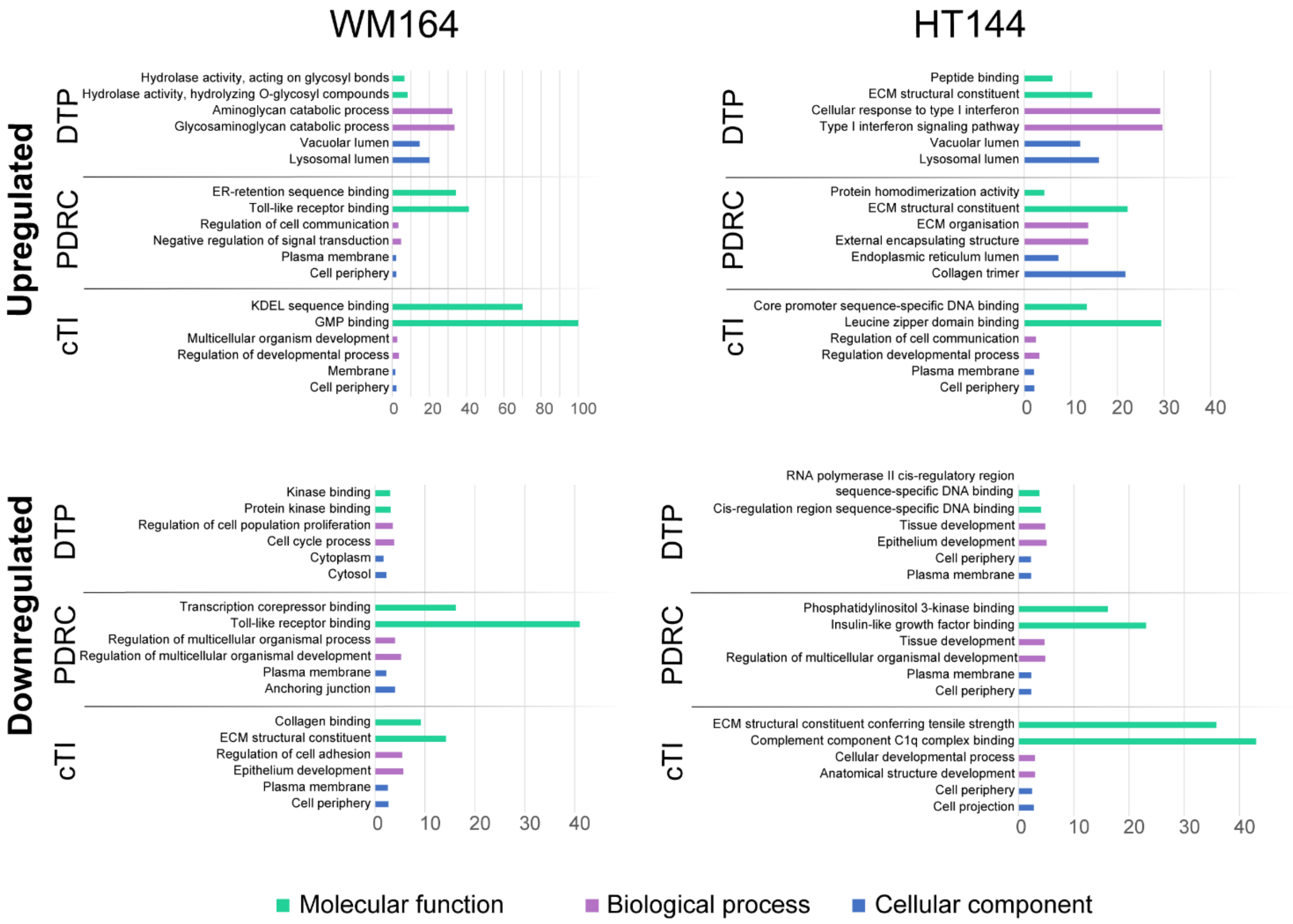


**Fig. S3 Gene ontology functional analysis of differentially expressed proteins in WM164 and HT144 treated with dabrafenib and cTI. Related to Fig. 3** Proteins from WM164 and HT144 cells treated with dabrafenib for 28 days (DTP) or for more than 105 days resistant (PDRC), and of cells treated with cTI for 13 weeks (cTI) were compared to proteins from untreated cells. Histograms represent fold enrichment of top 5 differentially expressed proteins ($p$-value ≤ 0.05; log2 fold change ± 1) involved in cellular component (blue), biological process (purple) and molecular function (teal) categories.

### Overexpression of proteins in DTP state that might relate to drug-resistance mechanisms, but not directly responsible for changes in cell membrane properties

Enzymes involved in fatty acid beta-oxidation (FAO) were also upregulated in dabrafenib-treated WM164 and HT144 cells (Fig. 3b), and even more so while in DTP state than in PDR state. This was previously observed with proteins found in the mitochondria and correlated with increased amounts of the metabolic intermediate and signal transducer acetyl-CoA and NADH [2]. Increased FAO was previously observed with DTPs of other BRAF mutant cell lines when exposed to MAPK inhibitors as a survival strategy before acquiring drug resistance [3, 4].

A number of lysosomal proteins are overexpressed in WM164 and HT144 DTPs, but not in PDRCs. In the DTP state, a total of 31 proteins within WM164 cells and 23 proteins within

HT144 cells (not shown in Fig. S2 as below top 10) had a fold enrichment of 15.4 and 11.3, respectively. In addition, proteins involved in the lysosomal lumen were also highly enriched in the cellular component analysis of both DTPs (Fig. S3). Overexpression of those enzymes is likely to increase entrapment of cytotoxic drugs in lysosomes, preventing them from reaching their intracellular target(s). Similarly, this could activate lysosome-associated signalling pathways, which could contribute to drug resistance [5, 6].

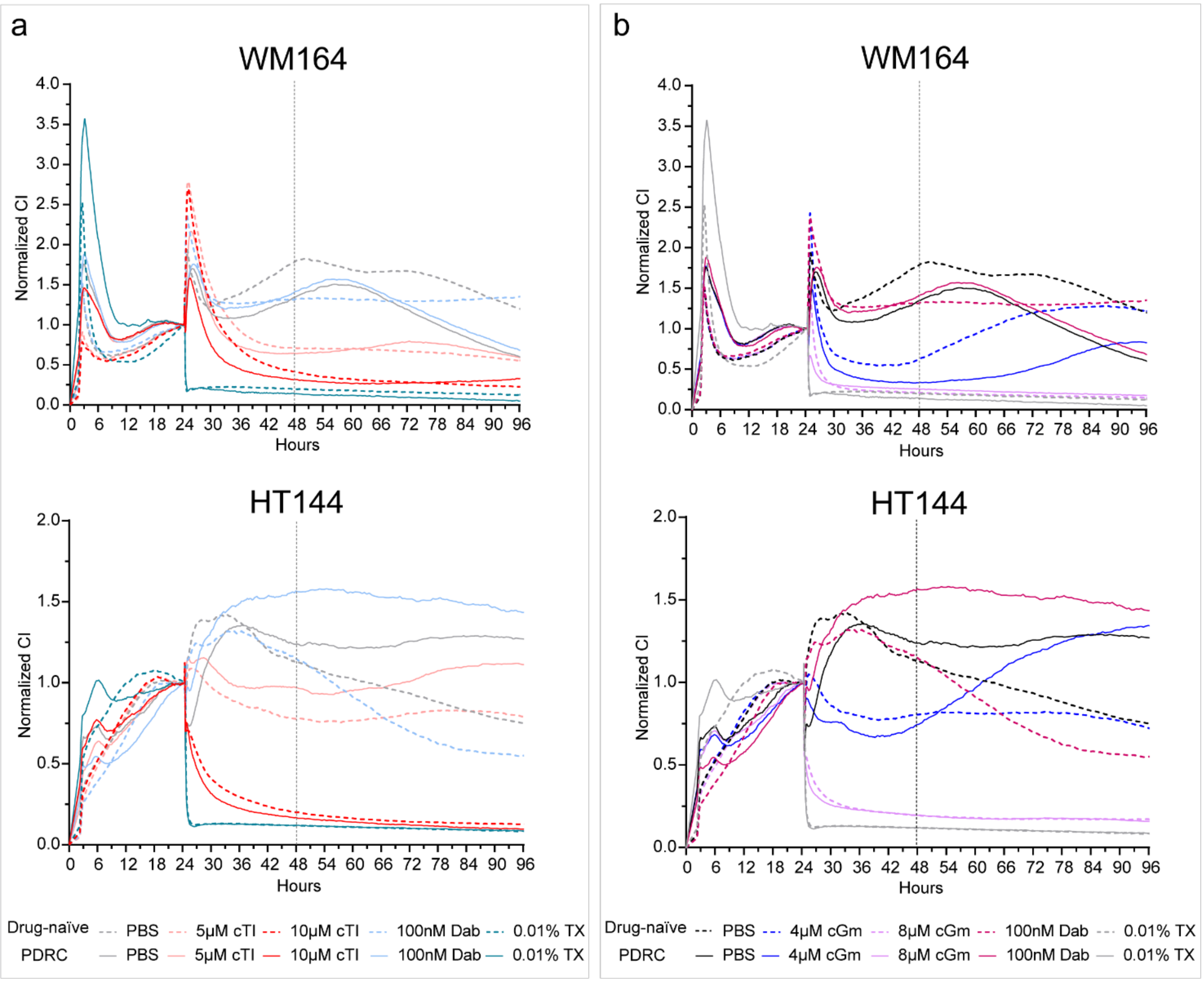


**Fig. S4 Proliferation of WM164 and HT144 cells when treated with cTI and cGm, monitored via cell impedance using XCELLigence**. XCELLigence traces of drug-naïve (dashed line) and drug-resistant (solid line) WM164 and HT144 cells incubated with cTI (**a**) and cGm (**b**). Cell viability correlates with cell index (CI) measured over time. 10,000 cells were seeded in each well of the E-plate 24 h before adding 100 nM dabrafenib or peptide. Samples were run in duplicate in three independent experiments. PBS control represents normal cell growth; 0.01% (v/v) triton X-100 (TX) induces cell death via cell membrane disruption, as confirmed with a CI close to 0. These results correlate with cytotoxicity results obtained with cells incubated with peptide for 24 h (see Fig. 4 and Fig. 1c).

**Co-treatments of melanoma cells with peptide and dabrafenib**

We quantified toxicity of cTI or cGm at fixed concentrations (equivalent to peptide $IC_{50}$), in the absence and presence of 50 nM dabrafenib to investigate whether co-treatment of peptide with small molecule BRAF-kinase inhibitors have additive or synergistic effect. Co-treatment of drug-naïve WM164 cells with cTI and dabrafenib for 72 h resulted in significantly higher toxicity (~80% cell death, instead of ~50% when the drug or peptide was added alone), suggesting an additive effect (Fig. S5, a). Co-treatment of PDRCs with either cTI or cGM, and dabrafenib did not increase toxicity compared to peptide alone (~50%), which agrees with these cells being resistant to dabrafenib and these two molecules having an additive effect.

The additive effect obtained with cTI can also be confirmed with dose response curve of dabrafenib co-treated with a fixed concentration of cGm or of cTI (Fig. S5, b). When increasing concentrations of dabrafenib were combined with 3 μM cTI, the percentage of dead cells for drug-naïve and drug-resistant WM164 was higher than cells incubated with cTI or dabrafenib alone.

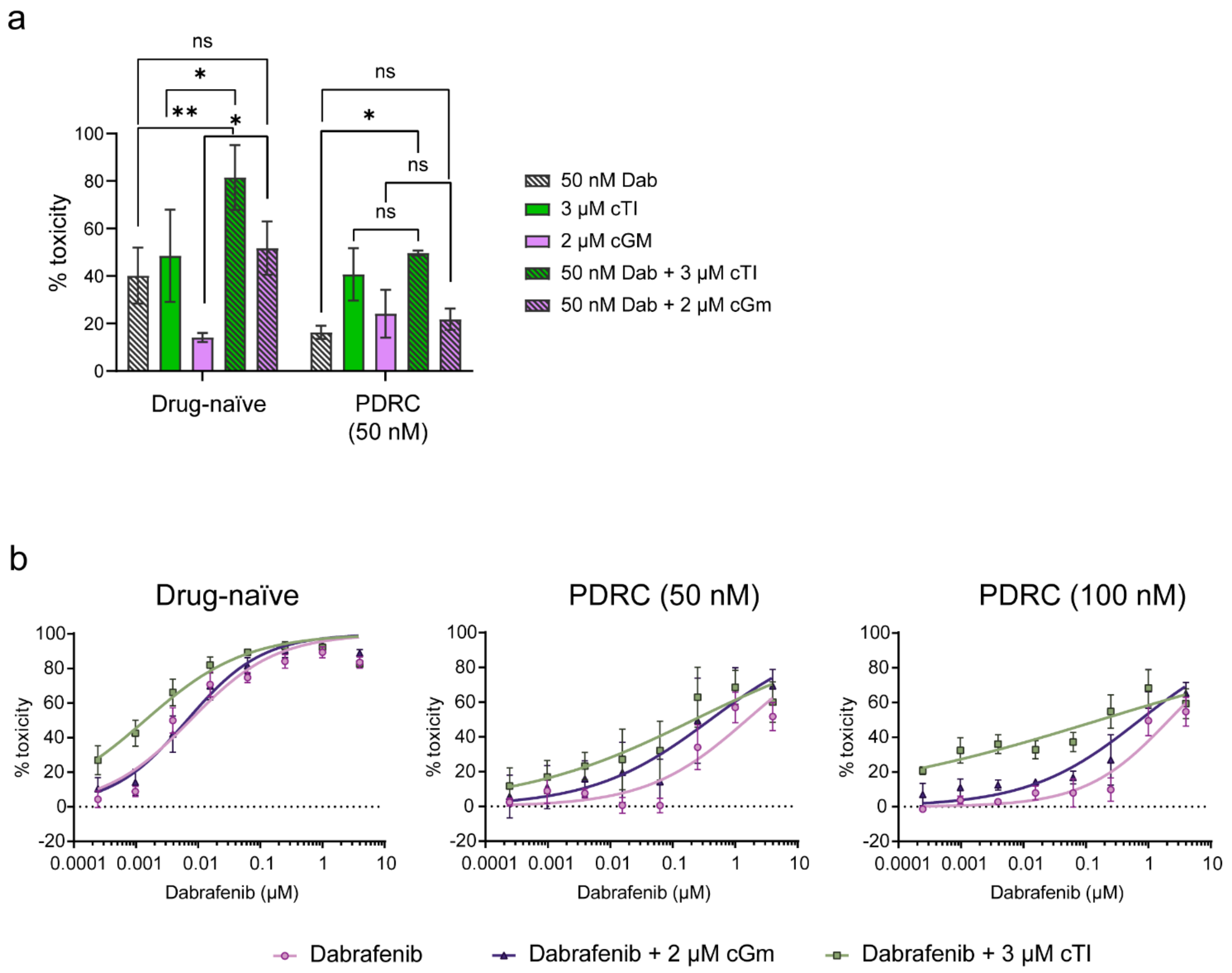


**Fig. S5 Cytotoxic effect of co-treatment of dabrafenib with peptide on drug-naïve and permanent drug resistant WM164 cells.** Fixed concentration for both peptides with 50mM dabrafenib (**a**) and with dose response curve of dabrafenib (**b**) are presented. Drug-naïve have not been incubated with dabrafenib and permanent drug-resistant cells were incubated for >100 days with 50 nM or 100 nM dabrafenib (PDRC). Resazurin assays were performed three days after addition of drug and peptides from three independent replicates each with cells generated from three independent experiments. Data points represent mean ± SD. Statistical difference between treatment determined by 2-way ANOVA with p-value <0.05 (*), ≤ 0.005 (**) and ≥ 0.05 (ns).

**Table S2** Mole ratio of glycerophospholipids (GPL), Sphingomyelin (SM) and cholesterol (Chol) during incubation of WM164 and HT144 with increasing concentration of cTI over 100 days. GPLs include PC, PE, PS and PI. Mean ratio ± SEM are shown (n=2). Related to Fig. 6c.

| | Days with treatment | % | | | SEM | | |
|---|---|---|---|---|---|---|---|
| | | GPL | SM | Chol | GPL | SM | Chol |
| WM164 | 0.5 µM cTI (2 d) | 73.5 | 3.2 | 23.4 | 0.1 | 0.1 | 0.2 |
| | 1 µM cTI (15 d) | 70.9 | 2.9 | 26.2 | 0.7 | 0.2 | 0.9 |
| | 2 µM cTI (60 d) | 73.8 | 3.6 | 22.5 | 1.2 | 0.2 | 1.0 |
| | 3 µM cTI (100 d) | 73.8 | 2.5 | 23.7 | 1.0 | 0.2 | 1.2 |
| | No peptide (15 d) | 72.2 | 2.9 | 24.9 | n/a | | |
| | No peptide (60 d) | 68.7 | 3.6 | 27.7 | | | |
| | No peptide (100 d) | 72.9 | 3.0 | 24.1 | | | |
| HT144 | 0.5 µM cTI (2 d) | 66.8 | 4.1 | 29.2 | 0.6 | 0.2 | 0.4 |
| | 1 µM cTI (15 d) | 70.2 | 3.4 | 26.4 | 0.4 | 0.0 | 0.4 |
| | 2 µM cTI (60 d) | 69.6 | 4.1 | 26.3 | 0.2 | 0.1 | 0.3 |
| | 3 µM cTI (100 d) | 70.9 | 3.0 | 26.2 | 0.2 | 0.1 | 0.1 |
| | No peptide (15 d) | 70.4 | 3.3 | 26.4 | n/a | | |
| | No peptide (60 d) | 68.0 | 3.8 | 28.2 | | | |
| | No peptide (100 d) | 72.1 | 3.3 | 24.6 | | | |

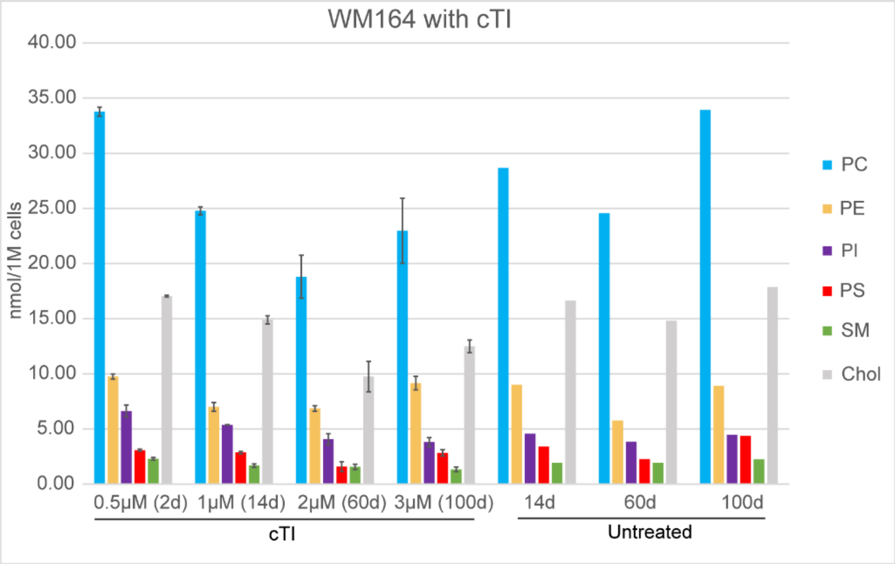


| | | cTI | | | | Untreated | | |
|---|---|---|---|---|---|---|---|---|
| | | 0.5μM (2d) | 1μM (14d) | 2μM (60d) | 3μM (100d) | 14d | 60d | 100d |
| nmol/1M cells | PC | 33.77 | 24.78 | 18.81 | 22.97 | 28.68 | 24.57 | 33.94 |
| | PE | 9.75 | 7.01 | 6.87 | 9.17 | 9.01 | 5.77 | 8.91 |
| | PI | 6.63 | 5.37 | 4.07 | 3.81 | 4.58 | 3.84 | 4.46 |
| | PS | 3.08 | 2.86 | 1.59 | 2.84 | 3.40 | 2.27 | 4.38 |
| | SM | 2.30 | 1.68 | 1.57 | 1.34 | 1.92 | 1.93 | 2.24 |
| | Chol | 17.05 | 14.89 | 9.75 | 12.49 | 16.63 | 14.81 | 17.88 |
| SEM | PC | 0.41 | 0.35 | 1.94 | 2.94 | | | |
| | PE | 0.23 | 0.39 | 0.25 | 0.61 | | | |
| | PI | 0.54 | 0.02 | 0.50 | 0.41 | | | |
| | PS | 0.09 | 0.12 | 0.43 | 0.30 | | | |
| | SM | 0.11 | 0.16 | 0.24 | 0.22 | | | |
| | Chol | 0.09 | 0.37 | 1.38 | 0.57 | | | |

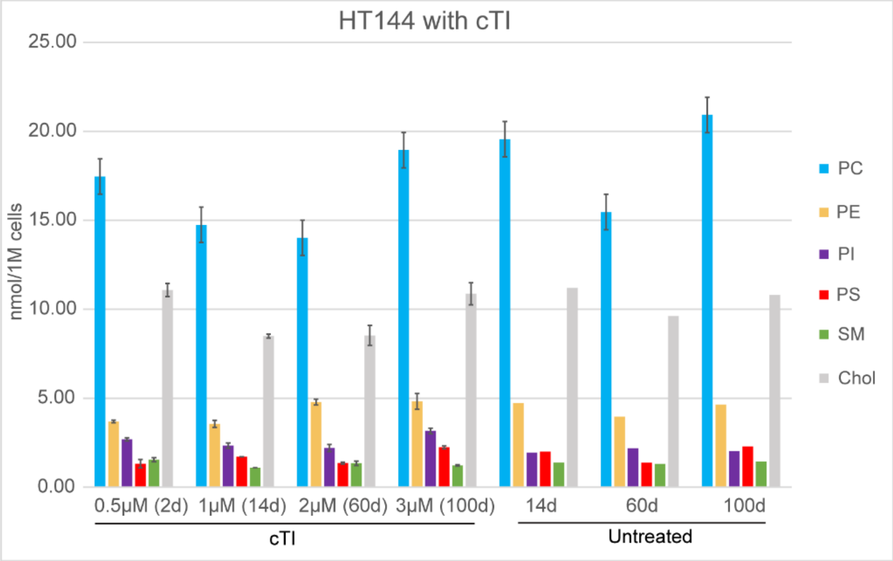


| | | cTI | | | | Untreated | | |
|---|---|---|---|---|---|---|---|---|
| | | 0.5μM (2d) | 1μM (14d) | 2μM (60d) | 3μM (100d) | 14d | 60d | 100d |
| nmol/1M cells | PC | 17.46 | 14.75 | 14.01 | 18.94 | 19.56 | 15.46 | 20.92 |
| | PE | 3.69 | 3.56 | 4.79 | 4.83 | 4.72 | 3.96 | 4.64 |
| | PI | 2.69 | 2.34 | 2.20 | 3.16 | 1.94 | 2.18 | 2.03 |
| | PS | 1.32 | 1.71 | 1.36 | 2.24 | 2.00 | 1.38 | 2.30 |
| | SM | 1.55 | 1.10 | 1.35 | 1.23 | 1.39 | 1.31 | 1.44 |
| | Chol | 11.08 | 8.49 | 8.53 | 10.87 | 11.20 | 9.61 | 10.81 |
| SEM | PC | 0.52 | 0.11 | 1.39 | 1.23 | | | |
| | PE | 0.07 | 0.20 | 0.16 | 0.44 | | | |
| | PI | 0.09 | 0.14 | 0.20 | 0.16 | | | |
| | PS | 0.24 | 0.00 | 0.05 | 0.08 | | | |
| | SM | 0.11 | 0.00 | 0.12 | 0.04 | | | |
| | Chol | 0.37 | 0.11 | 0.56 | 0.62 | | | |

**Fig. S6 Lipid composition of WM164 and HT144 cells while treated with cTI. Related to Fig. 6b and c.** Absolute quantification of each lipid class (PC, PE, PI, PS, SM and Chol) of WM164 and HT144 cells untreated and treated with 1 μM (15 days), 2 μM (60 days) and 3 μM (100 days) cTI in nmol per 1 million cells ± SEM (n = 2)
